# CRISPR-based screens uncover determinants of immunotherapy response in Multiple Myeloma

**DOI:** 10.1101/833707

**Authors:** Poornima Ramkumar, Anthony B. Abarientos, Ruilin Tian, Meghan Seyler, Jaime T. Leong, Merissa Chen, Priya Choudhry, Torsten Hechler, Nina Shah, Sandy W. Wong, Thomas G. Martin, Jeffrey L. Wolf, Kole T. Roybal, Andreas Pahl, Jack Taunton, Arun P. Wiita, Martin Kampmann

## Abstract

Cancer cells commonly develop resistance to immunotherapy by loss of antigen expression. Combinatorial treatments that increase levels of the target antigen on the surface of cancer cells have the potential to restore efficacy to immunotherapy. Here, we use our CRISPR interference and CRISPR activation-based functional genomics platform to systematically identify pathways controlling cell-surface expression of the multiple myeloma immunotherapy antigen - B cell maturation antigen, BCMA. We discovered that pharmacological inhibition of HDAC7 and the Sec61 complex increased cell-surface BCMA, including in primary patient cells. Importantly, pharmacological Sec61 inhibition enhanced the anti-myeloma efficacy of a BCMA-targeted antibody-drug conjugate. A CRISPR interference CAR-T coculture screen enabled us to identify both antigen-dependent and -independent mechanisms controlling response of myeloma cells to BCMA-targeted CAR-T cells. Thus, our study demonstrates the potential of CRISPR screens to uncover mechanisms controlling response of cancer cells to immunotherapy and to suggest potential combination therapies.

**Key Points:**

1. Using CRISPR screens, we systematically identify mechanisms increasing expression of the immunotherapy target BCMA and ADC efficacy.
2. We also identify antigen-independent mechanisms regulating response of cancer cells to BCMA-CAR-T cells.

## Introduction

Immunotherapy has transformed the treatment of many types of cancer including multiple myeloma (MM). B-cell maturation antigen (BCMA) is currently being evaluated in numerous clinical trials as an immunotherapy target in MM^1^. BCMA-targeted immunotherapy agents have shown improved responses in relapsed and refractory patients^2,3^. However, as with other MM therapies, resistance and relapse to BCMA-targeted therapies has emerged as a significant challenge and presents an unmet need^4,5^.

An important mechanism by which cancer cells can become resistant to different forms of immunotherapy in the clinic is the downregulation or loss of the targeted antigen^6, 7^, also termed “antigen escape”^6,8,9^. Ongoing clinical trials using BCMA-targeted chimeric antigen receptor T cells (CAR-T cells) have reported antigen loss in some patients undergoing relapse^4,5^, indicating that reduced cell-surface levels of BCMA may be an important mechanism of therapy resistance. However, the underlying cellular mechanisms remain to be understood. CRISPR-based genetic screens are a powerful research tool to define mechanisms of treatment resistance in cancer cells to different immunotherapies^10–12^, to design strategies to overcome resistance^13 11^, identify novel immunotherapy target antigens^14^ and better understand immune checkpoint regulation^13^.

In this study we used our CRISPR-interference/CRISPR-activation (CRISPRi/CRISPRa) functional genomics platform^15,16^ to systematically elucidate the mechanisms by which the cellsurface expression of BCMA is controlled in MM cells, and to test whether some of these mechanisms would be potential targets for combination therapy to enhance BCMA-directed immunotherapy. Furthermore, we conducted a CRISPRi screen for genes controlling sensitivity of MM cells to BCMA-directed CAR-T cells. To our knowledge, this is the first genetic screen for genes in multiple myeloma controlling response to CAR-T cells directed against a clinically relevant target. Our results demonstrate the potential of CRISPR screens to elucidate mechanisms controlling the response of cancer cells to immunotherapy and the identification of potential pharmacological strategies to enhance immunotherapy.

## Materials and Methods

### CRISPRi and CRISPRa flow cytometry screen

CRISPRi and CRISPRa cell lines were generated as detailed in the supplemental materials and methods. For transduction of each sublibrary, AMO1 cells expressing the CRISPRi or CRISPRa machinery were spin-infected with the virus at 700 g for 2 hrs at 32°C. 48 hrs later the cells were analyzed for percentage of infection by flow cytometry and treated with 1 μg/ml of puromycin to obtain a pure population of sgRNA expressing cells. On day 12 and day 5 post infection with the CRISPRi and CRISPRa sublibraries, cells were stained for cell surface BCMA and flow sorted to enrich for populations of cells expressing low or high cell surface levels of BCMA. Briefly, for each sublibrary, cells were resuspended in FACS buffer (PBS containing 0.5% FBS) at a concentration of 10*10^6^ cells/ml. The cells were blocked using Human BD Fc Block (#564220; BD Biosciences); stained with PE/CY7-BCMA (19F2) (#357508, Biolegend) antibody and resuspended in FACS buffer for flow sorting. Top and bottom 30% of cells expressing BCMA as determined from PE/Cy7-BCMA histogram were flow sorted using BD FACS Aria II. The different cell populations were then processed for next-generation sequencing as described previously ^15,17^ and sequenced on an Illumina HiSeq-4000. To identify significant hit genes, sequencing reads were analyzed using the MAGeCK-iNC pipeline as described previously^18^.

### CRISPRi validation screen

sgRNAs targeting the selected 41 top hits identified from the primary CRISPRi screen were cloned into a custom library of 90 sgRNAs, including two sgRNAs per gene and 8 non-targeting control sgRNAs. The sgRNAs were transduced into a panel of CRISPRi-MM cell lines (KMS11, AMO1, RPMI8226, OPM2 and KMS12-PE). The validation screen was carried out similar to the primary screen where cells were stained using PE/Cy7-BCMA or FITC-CD38 (#303504, Biolegend). Knockdown phenotypes for both BCMA and CD38 were hierarchically clustered based on Pearson correlation using Cluster 3.0^19^ and Java TreeView 3.0 (http://jtreeview.sourceforge.net/)^20^.

### Antibody drug conjugate dose response assays

BCMA-targeted antibody drug conjugate (ADC), HDP-101 was produced as described previously^21,22^, For drug combination studies, cells were treated with either DMSO or indicated concentration of drugs prior to seeding. Increasing concentrations of the HDP-101 were then added to cells to obtain a dose response curve. 96 hrs post treatment, cell viability was measured using CellTiter-Glo 2.0 reagent (#G9241, Promega) following manufacturer’s protocol. Raw luminescence signals were collected using a SpectraMax M5 plate reader (Molecular Devices). All measurements were taken in triplicates and raw counts were normalized as the percent of signal relative to untreated cells or percent maximum signal when comparing more than one drug. Sigmoidal dose-response curve fitting for IC50 calculation was performed using Prism V7 package (GraphPad software Inc.).

### CAR-T cell cytotoxicity assay

Generation of CAR-T cells is detailed in the supplemental materials and methods. For T cell cytotoxicity assays, GFP-CAR-T cells^23^ were co-cultured in the presence of BFP-CRISPRi MM cells stained with eFluor-670 (#65-0840-85, Thermo Fisher) at a ratio of 1:1 for 24 hrs. Baseline target cell death was measured by incubating CRISPRi MM cells without any CAR-T cells over the same time period. Cells were then stained with BV786-CD69 (FN50) (#310932, Biolegend) to determine T cell activation status and propidium iodide to assess overall cell death.

### CRISPRi-CAR survival screen

AMO1 CRISPRi cells expressing an sgRNA library targeting 12,838 genes (including kinases, phosphatases, cancer drug targets, apoptosis genes, proteostasis genes and mitochondrial genes) were co-cultured in the presence or absence of GFP-BCMA CAR-T cells at a ratio of 1:1 for 24 hrs. Surviving cells were then harvested and the different cell populations were processed for next generation sequencing as described previously^15,17^. Sequencing reads were analyzed using MAGeCK-iNC pipeline developed in our lab to identify significant hit genes as described^18^.

## Results

### Genome-wide CRISPR screens to identify genes controlling cell surface BCMA expression

To identify novel genes or pathways regulating cell surface expression of BCMA in MM cell line, we performed genome-wide CRISPRi and CRISPRa screens in a MM cell line (Fig 1A). Performing parallel CRISPRi knockdown and CRISPRa overexpression screens can yield complementary insights^15,24^. Knockdown of individual genes by CRISPRi can inhibit the function or activity of an entire pathway or protein complexes for which the individual gene is required – whereas overexpression of an individual gene does not in general enhance the function of an entire pathway or protein complex. However, CRISPRi screens can only interrogate the function of genes expressed in the cell line used in the screen, whereas CRISPRa can uncover consequences of inducing a gene that is not normally expressed.

**Fig 1:**
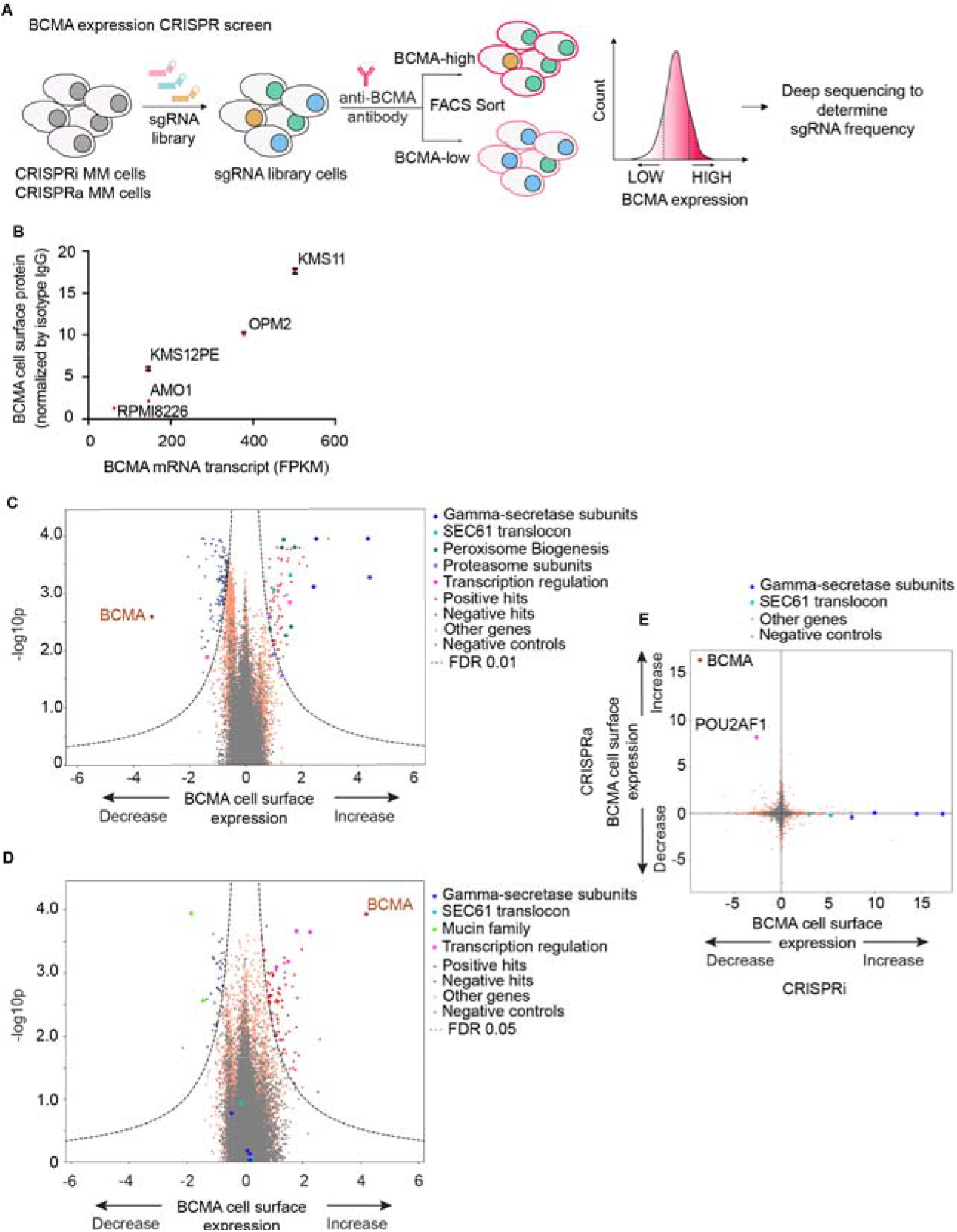
Genome-wide CRISPRi/a screens to identify genes regulating cell-surface expression of BCMA. **A,** Schematic representation of our genome-wide CRISPRi and CRISPRa screens to identify modulators of BCMA expression. AMO1 cells constitutively expressing the CRISPRi or CRISPRa machinery were transduced with genome-wide lentiviral sgRNA library. Following transduction, cells were stained for cell surface levels of BCMA and sorted by fluorescence-activated cell sorting (FACS) to enrich for populations with low or high levels of cell surface BCMA. Frequencies of cells expressing a given sgRNA were determined in each population by next generation sequencing. **B,** BCMA expression levels in a panel of MM cell lines. BCMA transcript FPKM levels obtained from the Keats lab database: https://www.keatslab.org were plotted against cell surface expression levels of BCMA quantified by flow cytometry. The flow cytometry data are means of three biological replicates, error bars denote SD. Note that some error bars are not visible because values are small. **C and D,** Volcano plots indicating the BCMA expression phenotype and statistical significance for knockdown (CRISPRi, C) or overexpression (CRISPRa, D) of human genes (orange dots) and quasi-genes generated from negative control sgRNA (grey dots). Hits genes corresponding to functional categories are color-coded as labeled in the panel. **E,** Comparison of phenotypes from the CRISPRi and CRISPRa screens. Selected hit genes are color-coded.

To select a suitable cell line for these primary screens, we compared levels of BCMA expression on the RNA (www.keatslab.org) and cell surface protein levels for a panel of MM cell lines (Fig. 1B). We found that AMO1 cells expressed moderate levels of cell surface BCMA. We therefore selected this line for our primary screen as we reasoned it would enable us to identify modifiers that either increase or decrease BCMA surface levels (Fig 1B).

AMO1 cells were lentivirally transduced to express the CRISPRi and CRISPRa machinery and CRISPR functionality was tested using sgRNA targeted towards *CD38* and *CXCR4* respectively (Supplemental Fig. 1). The genome-wide screen was conducted as shown in Fig 1A. Briefly, AMO1 cells expressing the CRISPRi and CRISPRa machinery were transduced with pooled genome-wide sgRNA library. The cells were then stained for cell surface BCMA using fluorescent tagged antibody and subjected to flow sorting into BCMA-low and BCMA-high populations. Frequencies of cells expressing each sgRNA were identified by nextgeneration sequencing. The genome-wide CRISPRi screen identified several genes regulating cell surface expression of BCMA (Fig. 1C and Supplemental Table 1). Knocking down BCMA itself or its transcription factor *POU2AF1*^25^ both resulted in significant downregulation of cell surface BCMA expression, thus validating the screen (Fig 1C). Furthermore, all of the subunits of the gamma secretase complex were among the top hits, and their knockdown resulted in a significant increase in cell surface BCMA (Fig 1C). This was consistent with previous reports showing that gamma secretase complex cleaves membrane-bound BCMA (mBCMA) to a soluble form (sBCMA)^9,26^. Moreover, we identified several functional categories of genes regulating expression levels of BCMA including subunits of the Sec61 translocon complex, peroxisome biogenesis, proteasome subunits, and regulators of transcription (Fig 1C).

Our CRISPRa genome-wide screen (Fig. 1D and Supplemental Table 2) identified genes in the Mucin family (*MUC1, MUC21*) and several genes involved in transcriptional regulation (*POU2AF1, CBFA2T3, MAML2, RUNX3*) that regulated surface BCMA. Systematic comparison of the parallel CRISPRi and CRISPRa screens (Fig. 1E) revealed that overexpression and knockdown of some genes had opposing effects on BCMA surface levels – including BCMA itself and *POU2AF1*. However, other genes were hits in only the CRISPRi or CRISPRa screens, notably the subunits of the gamma secretase complex and the Sec61 complex; highlighting the fact that CRISPRi and CRISPRa screens can uncover complementary results, as we previously _described_^15,24^.

### Validation of hit genes in a panel of multiple myeloma cell lines

To test the generality of our findings, we decided to validate hit genes from the primary screen performed on the AMO1 cells in a panel of CRISPRi-MM lines, in which we confirmed CRISPRi activity (Supplemental Fig. 1). We chose 41 hits both based on strength of the phenotype from the CRISPRi screen, and based on their potential as therapeutic targets. The secondary screen was performed similarly to the genome-wide screen to validate changes in BCMA. The screen was carried out in parallel for a different cell surface protein expressed in all MM lines, CD38, to investigate whether hit genes selectively affected BCMA expression.

As observed in the primary CRISPRi screen, knockdown of the subunits of gamma secretase subunits significantly increased cell surface BCMA (Fig. 2A). We found this effect to be selective for BCMA, since gamma secretase knockdown had no effect on CD38 expression in a panel of MM cell lines (Fig 2A). This was further validated using a pharmacological inhibitor of gamma secretase complex, RO4929097^27^. MM cell lines treated with the indicated concentration of RO4929097 showed an increase in both BCMA cell surface expression by flow cytometry (Fig 2B) and in total protein levels of BCMA by immunoblotting (Fig 2C). Thus, we were able to successfully validate genes from our primary screen in a panel of MM cell lines, using both genetic and pharmacological perturbations.

**Fig 2:**
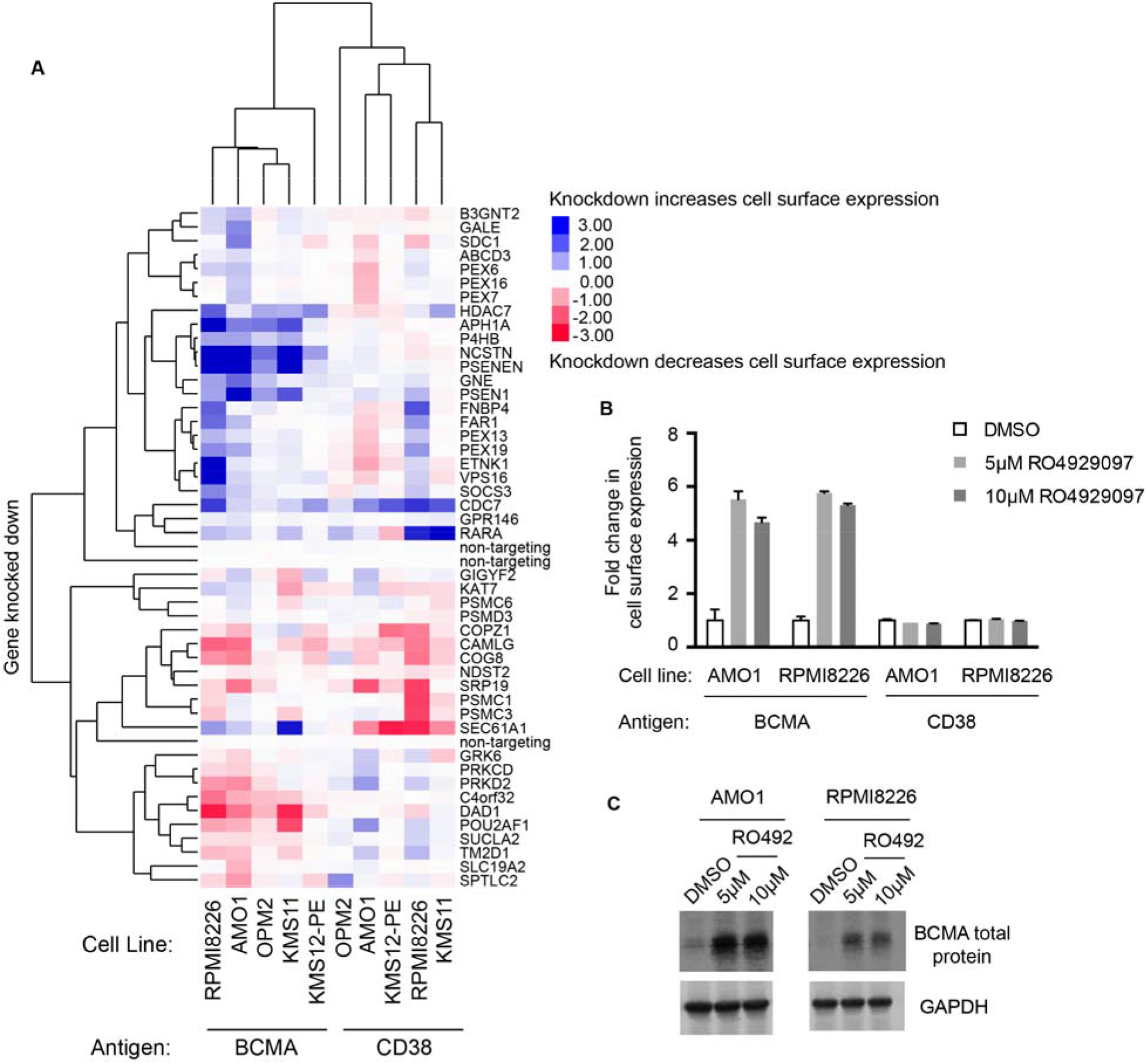
Validation of hit genes from the primary screen. **A**, Heat map representation of knockdown phenotype scores from CRISPRi validation screens in a panel of MM cell lines, for cell-surface levels of BCMA and CD38. Both genes and screens were hierarchically clustered based on Pearson correlation. B and C, AMO1 and RPMI8226 cells treated with indicated concentrations of the gamma-secretase inhibitor RO4929097 or DMSO for 48 hrs were analyzed by flow cytometry (B) and immunoblotting (C) for changes in BCMA levels. (B) Fold changes in BCMA cell surface levels were determined by normalizing to DMSO-treated cells. Data points are means of three biological replicates; error bars denote SD. (C) Western blot of endogenous BCMA. GAPDH was used as a loading control.

Several of the novel factors from our primary screen also validated in the larger panel of MM cells (Fig. 2A and Supplemental Tables 3,4), most notably the transcriptional regulator *HDAC7* and *SEC61A1*, a subunit of the SEC61 translocon. Knockdown of both factors increased cell-surface levels of BCMA, but not CD38 (Fig 2A).

### Class IIa HDAC inhibition upregulates BCMA transcription

Our primary CRISPRi screen showed that knockdown of *HDAC7*, but not any other HDAC gene, upregulates BCMA expression (Fig. 3A). *HDAC7* is a member of the histone deacetylase^28^ (HDAC) family, which have emerged as crucial transcriptional co-repressors. *HDAC7* belongs to the Class II family of HDACs, which are subdivided into Class IIa including *HDAC4,5,7,9* and Class IIb including *HDAC6* and *HDAC10*^28^. We validated our screen findings using a pharmacological inhibitor targeting Class IIa HDACs, TMP269^29^. Treatment of RPMI8226 cells with increasing concentrations of TMP269 showed a two-fold increase in BCMA protein levels (Fig 3B-D). Because HDACs have been established to regulate transcription of several genes, we performed qPCR to analyze for changes in transcript levels of BCMA. Our results showed a two-fold increase in BCMA transcript levels (Fig 3E), indicating that Class II HDAC inhibition regulates transcription of BCMA. No significant change was observed in the CD38 protein or transcript levels (Fig 3B, 3E). Furthermore, treatment of RPMI8226 cells with a combination of TMP269 and gamma-secretase inhibitor, RO4929097 showed a further increase in BCMA surface expression levels, supporting the notion that they act through different mechanisms (Fig 3F).

**Fig 3:**
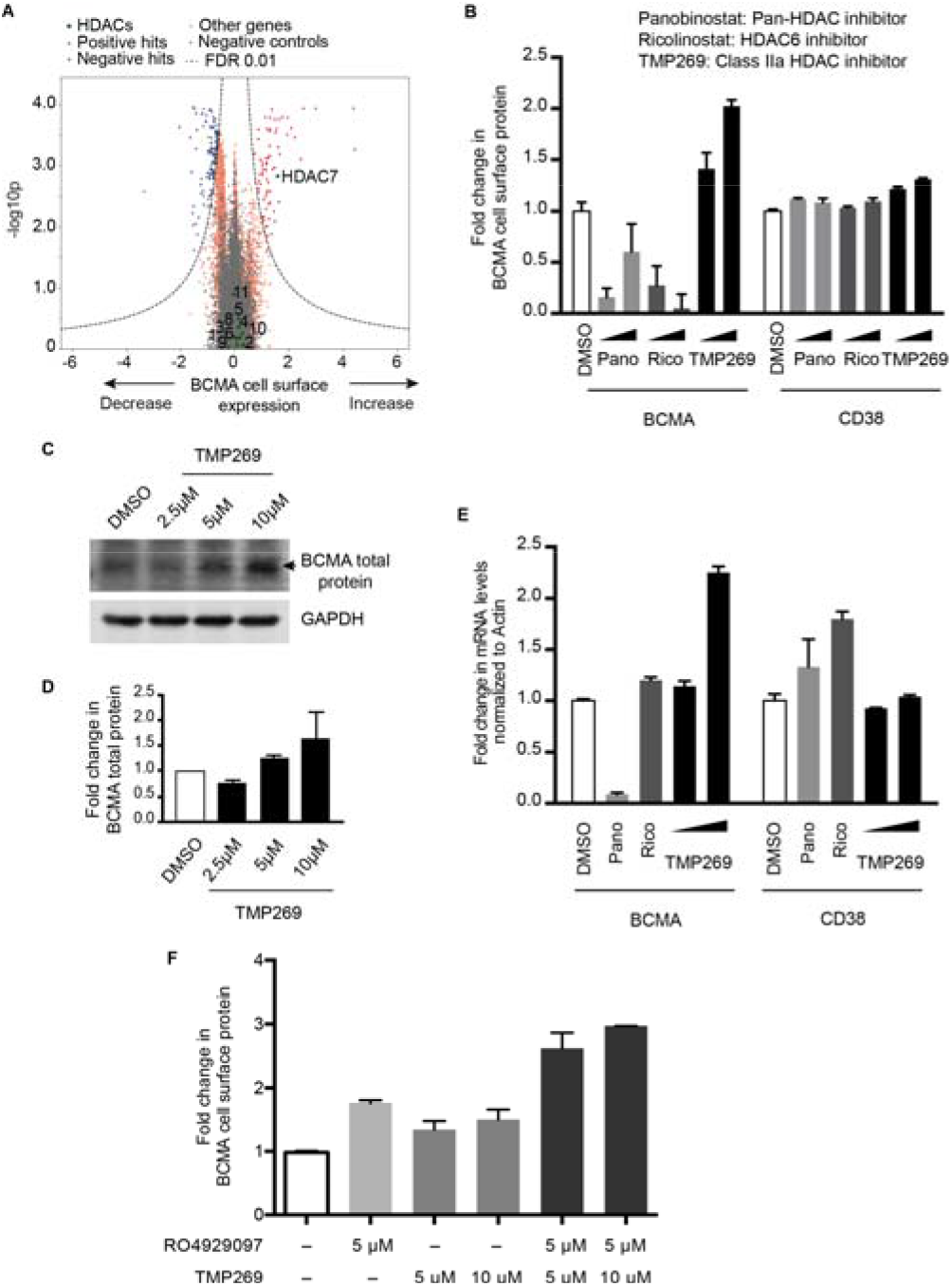
Class IIa-HDAC inhibition increases transcription of BCMA. **A,** Volcano plot indicating the BCMA expression phenotype and statistical significance for knockdown (CRISPRi) of human genes (orange dots) and quasi-genes generated from negative control sgRNA (grey dots). Genes in the HDAC family are shown as green dots and labeled with the HDAC number. **B,** RPMI8226 cells were treated with increasing concentrations of the pan HDAC inhibitor panobinostat (10 nM, 25 nM); the HDAC6-specific inhibitor ricolinostat (0.5 μM, 1 μM); the class II-HDAC inhibitor TMP269 (5 μM, 10 μM), or DMSO for 48 hrs and analyzed by flow cytometry for cell surface expression of BCMA and CD38. Fold changes in protein levels were determined by normalizing to the DMSO-treated cells. Data points are means of three biological replicates; error bars denote SD. **C and D,** Total protein extracts from RPMI8226 cells treated with 2.5 μM, 5 μM and 10 μM of TMP269 for 48 hrs were analyzed by immunoblotting for expression levels of BCMA. GAPDH was used to normalize differences in loading amounts. Data is represented as fold change relative to the total protein expression level after normalization with GAPDH. Data points are means of two technical replicates, error bars denote SD. **E,** RPMI8226 cells treated with 10 nM panobinostat, 0.5 μM ricolinostat and 5 μM and 10 μM of TMP269 for 48 hrs were processed for quantitative PCR (qPCR) to determine transcript levels of BCMA and CD38. Fold changes in transcript levels with different drug treatments were determined after normalizing to beta-actin gene. Data are means of two biological replicates, error bars denote SD. **F,** RPMI8226 cells were treated with DMSO; 5 μM and 10 μM of TMP269; 5 μM of gamma-secretase inhibitor RO4929097 as single agents or in combination for 48 hrs and analyzed by flow cytometry for cell surface expression of BCMA. Fold changes in protein levels were determined by normalizing to the DMSO-treated cells. Data points are means of three biological replicates; error bars denote SD.

To investigate whether this change in BCMA levels is specific to HDAC7 inhibition, we treated cells with a pan-HDAC inhibitor, panobinostat^30^, and a HDAC6-specific inhibitor, ricolinostat^31^ Our results showed no increase in BCMA transcript levels with these agents (Fig.3B,E). In fact, these compounds resulted in a decrease in BCMA cell surface levels, which can be rationalized by three facts. First, while panobinostat is a pan-HDAC inhibitor, it is not very potent for HDAC7, and we are using it at nanomolar concentrations well below the IC50 for HDAC7 (1.378 μM^32^), since we are limited by toxicity. Second, panobinostat inhibits several different HDACs with a range of different cellular functions, and will therefore cause pleiotropic effects. Second, Both panobinostat and ricolinostat are very toxic to MM cells, resulting likely in a range of cellular changes due to toxicity, whereas we found no toxicity in MM cells with TMP269 treatment (Supplemental Fig. 2A).

Furthermore, treatment with TMP269 in K562 cells, which do not express BCMA, did not lead to an increase in BCMA expression (Supplemental Fig. 2B,C). These results indicate the potential for using Class IIa HDAC inhibition to increase expression of BCMA in plasma cells in the context of BCMA-targeted immunotherapy.

### The Sec61 translocon regulates BCMA protein levels

Most integral plasma membrane proteins are inserted into membranes via the Sec61 translocon, located in the endoplasmic reticulum. It has been shown that inhibition of Sec61 affects correct localization of a subset of membrane proteins^33,34^. Surprisingly, our CRISPRi screen identified that knockdown of genes in the SEC61 pathway, such as the *SEC61A1, SEC61G, SSR1* and *OSTC*, resulted in an increase – rather than a decrease - in BCMA cell surface levels (Fig. 1C). This unexpected finding was confirmed in our validation screens in the panel of MM cell lines (Fig. 2A). Conversely, *SEC61A1* knockdown resulted in a decrease in CD38 cell surface levels (Fig. 2A), as would be expected for most membrane proteins.

To test whether pharmacological inhibition of the Sec61 translocon complex phenocopies the genetic knockdown, we treated cells with SEC61 inhibitors, CT8^33^ and PS3061^35^. We observed that treatment of MM cell lines with increasing concentrations of these compounds results in an up to five-fold dose-dependent increase in cell-surface BCMA levels and a decrease in CD38 levels with minimal cytotoxicity, as evidenced by flow cytometry (Figs. 4A, 4D, Supplemental Fig. 3). Moreover, immunoblotting also showed a dose-dependent increase in total BCMA protein levels (Fig. 4B, 4E, Supplemental Fig 3) and up to two-fold increase in BCMA transcript levels (Fig. 4C, 4F). Furthermore, treatment of cells with SEC61 inhibitors resulted in a decrease in TACI, which, like BCMA, belongs to the TNFRSF family of proteins (Supplemental Fig 3), indicating that the SEC61 inhibition selectively upregulates BCMA levels. Combinatorial treatment with both gamma-secretase and SEC61 inhibitors had cell-line specific outcomes, highlighting the heterogeneity of MM cell lines. We observed an increase in BCMA expression when both gamma-secretase and SEC61 inhibitors were combined in AMO1 cells (Fig. 4G), but not in KMS11 cells (Fig. 6D).

**Fig 4:**
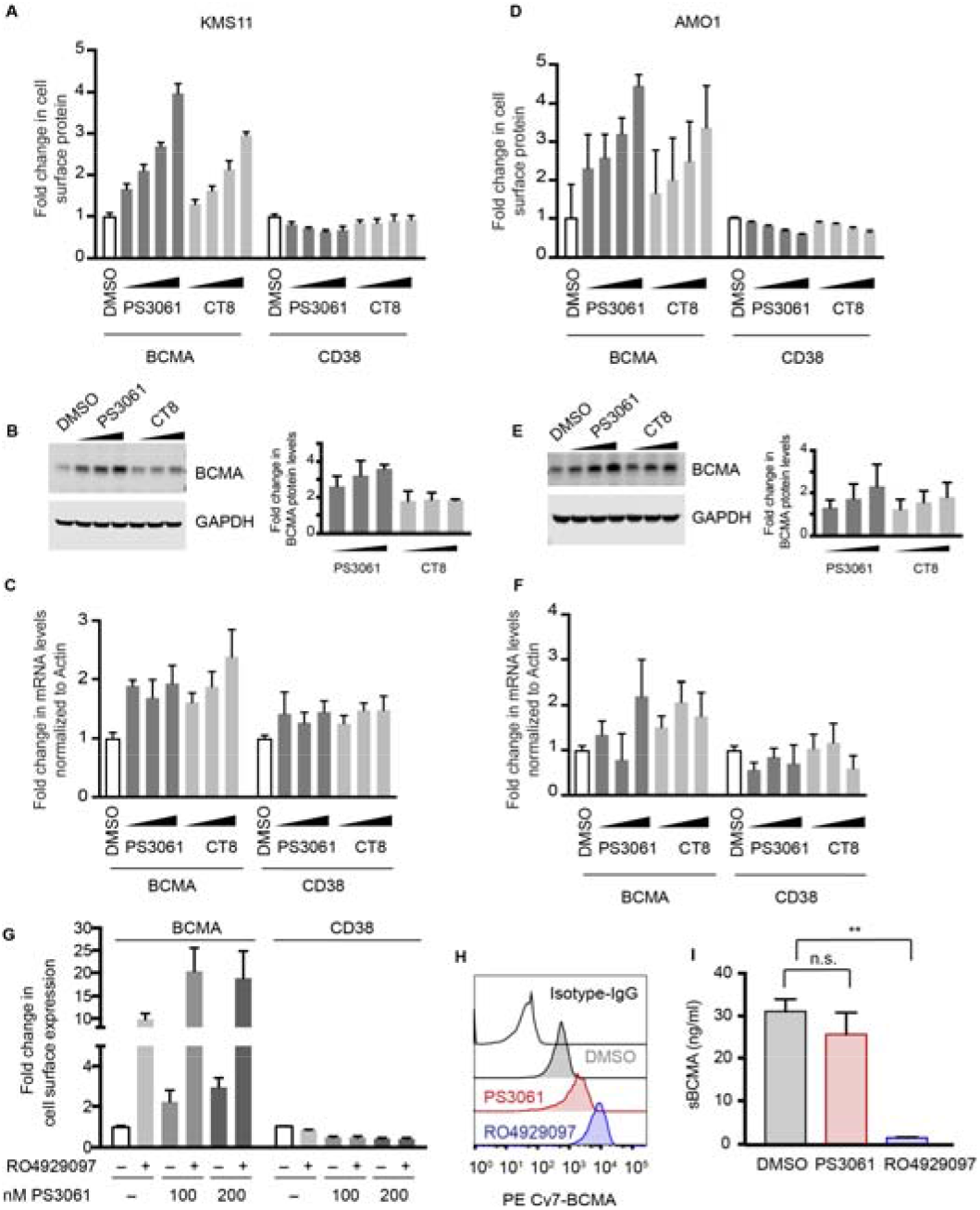
Sec61 inhibitors increase cell-surface expression of BCMA in KMS11. **(A-C) and AMO1 (D-F) cells A and D,** KMS11 and AMO1 cells were treated with increasing concentrations of Sec61 inhibitors, CT8 and PS3061 (100, 200, 400, 800 nM) or DMSO as a control for 24 hrs. Cells were stained for cell surface expression of BCMA and CD38 and analyzed by flow cytometry. Data are means of three biological replicates; error bars denote SD. **B and E,** Total protein lysates from cells treated with increasing concentration of CT8 and PS3061 (200, 400, 800 nM) for 24 hrs were processed for western blotting. GAPDH was used to normalize for differences in loading amounts. Data is represented as fold change relative to the normal protein expression level after normalization with GAPDH. Data points are means of two technical replicates, error bars denote SD. **C and F,** cells treated with increasing concentrations of PS3061 and CT8 (200, 400, 800 nM) for 24 hrs were processed for qPCR to determine transcript levels of BCMA and CD38. Fold change in transcript levels were determined after normalizing to beta-actin. Data are means of two biological replicates, error bars denote SD. **G,** AMO1 cells were treated for 24 h with the Sec61 inhibitor, PS3061 (100, 200 nM) and the gamma-secretase inhibitor RO4929097 (5 μM) as single agents or in combination using DMSO as a control. Cells were stained for cell surface expression of BCMA and CD38 and analyzed by flow cytometry. Data are means of three biological replicates; error bars denote SD. **H and I,** KMS11 cells were treated with 800 nM PS3061, 5 μM RO4929097 and DMSO for 24 hrs. G, Drug-treated cells were analyzed by flow cytometry for cell surface expression of BCMA. Histograms indicate distribution of PE/CY7 BCMA in the drug-treated cells. Data is a representation of two biological replicates. H, Soluble BCMA (sBCMA) concentration in the cell culture supernatant post drug treatment was measured using ELISA. Data are means of two biological replicates, error bars denote SD. \**P* < 0.005, n.s. - not significant, two-tailed unpaired t test.

To uncover the mechanism by which SEC61 inhibition increases BCMA levels, we first tested the possibility that a reduction in gamma-secretase levels at the plasma membrane would reduce shedding of BCMA from the cell surface. Pharmacological inhibition of either the gamma-secretase or Sec61 resulted in increased mBCMA as evidenced by flow cytometry (Fig. 4H). To determine levels of sBCMA generated by gamma-secretase processing, we performed sandwich ELISA assay on cells treated with PS3061 or RO4929097. Our results show that inhibition of gamma-secretase activity significantly reduced sBCMA levels, whereas we did not observe significant changes in sBCMA levels with SEC61 inhibition (Fig 4I). This finding indicated that the increase in cell-surface BCMA is not driven by reduced shedding of BCMA.

To validate our findings in primary patient cells, we treated bone marrow mono-nuclear cells derived from different MM patients (Supplemental Table 5) with increasing concentrations of TMP269 and RO4929097 for 24 hrs and analyzed the expression of BCMA on plasma cells by flow cytometry. We observed an up to 2.5-fold increase in cell-surface BCMA with class IIa HDAC inhibition and an up to 8-fold increase with gamma-secretase inhibition (Fig. 5). Similarly, treatment with the Sec61 inhibitor PS3061 increased cell-surface levels of BCMA in primary patient cells approximately two-fold, while not affecting cell-surface levels of CD38 (Fig. 5). Furthermore, moderate effects on BCMA expression was observed on non-plasma cells in the patient samples (Supplemental Fig. 4).

**Fig 5:**
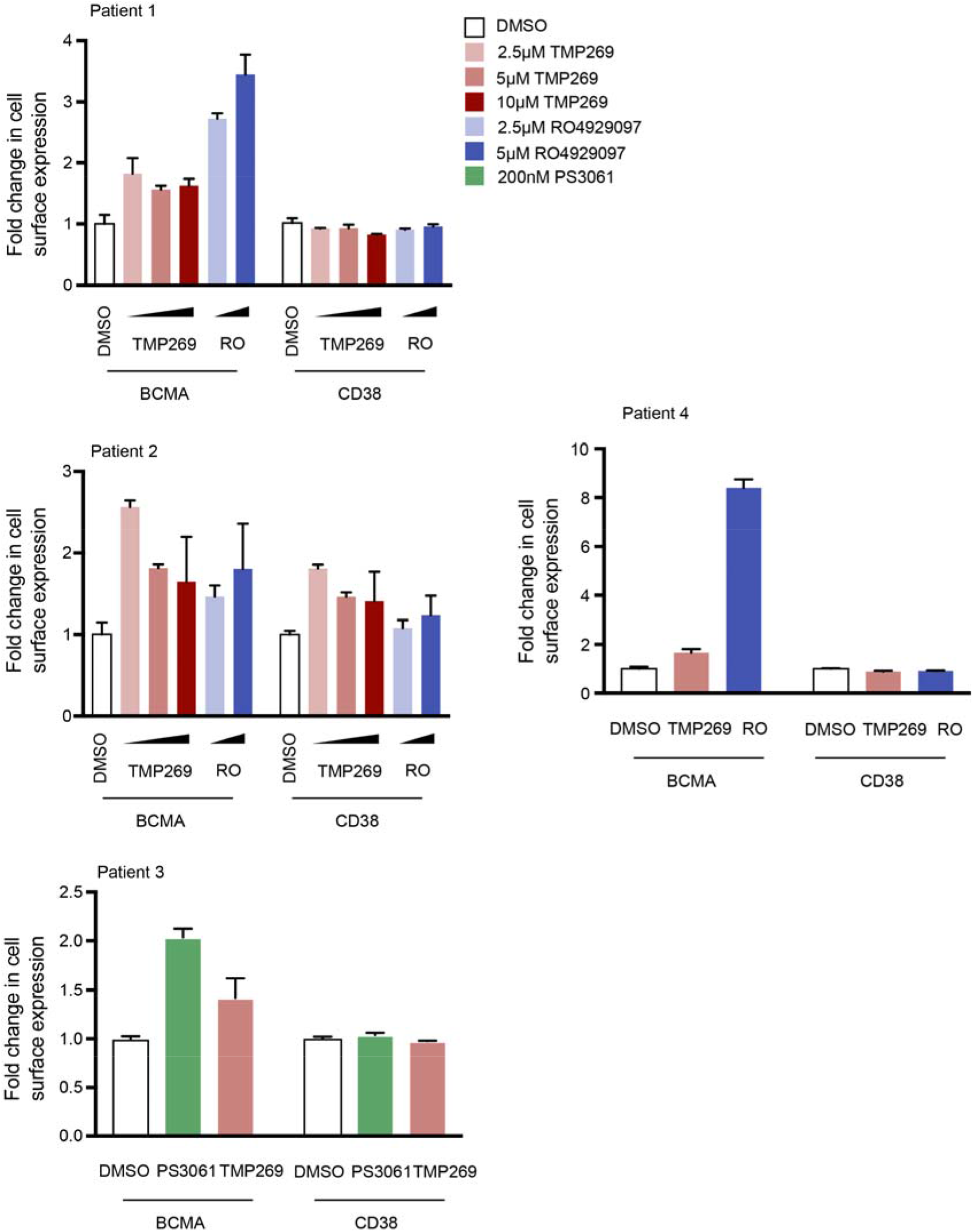
Pharmacological inhibition of the validated hits in myeloma patient samples upregulate cell-surface BCMA levels. Bone marrow-mononuclear cells (BM-MNCs) isolated from bone marrow aspirates from different MM patients were treated with indicated concentration of Class II-HDAC inhibitor, TMP269; gamma-secretase inhibitor, RO4929097; and Sec61 inhibitor, PS3061 for 24 hrs. Cells were stained for cell-surface CD138, BCMA and CD38 and analyzed by flow cytometry. Fold change in BCMA and CD38 levels were determined in CD138+ live cell populations. Data are means of three technical replicates, error bars denote SD.

### Increased efficacy of BCMA-ADC when combined with Sec61 inhibitor

Immunotherapy agents such as ADC are sensitive to changes in expression of target antigen ^36^. We obtained an ADC targeting BCMA, HDP101^21,22,37^ developed by Heidelberg Pharma that has proven efficacy in MM cell lines at sub-nanomolar concentrations (unpublished data). Here, we knocked down BCMA using two independent sgRNAs in CRISPRi expressing KMS11 cells (Supplemental Fig. 5) and performed a dose-response assay using increasing concentrations of BCMA-ADC. Our results indicate that cell lines in which BCMA is knocked down no longer respond to the ADC, thus establishing that the therapy is dependent on expression levels of BCMA on the cell surface (Fig 6A). Moreover, when cells were treated with either the gamma secretase inhibitor RO4929097 or the Sec61 inhibitor PS3061 at concentrations increasing cell surface levels of BCMA (Fig. 6B) in combination with HDP101, we observed increased efficacy of the BCMA-ADC on the MM cell lines (Fig 6C). Furthermore, we tested the effects of combining gamma-secretase and Sec61 inhibition on sensitivity to HDP101 in KMS11 cells. We did not observe a further sensitization of the cells to the ADC when the inhibitors are combined (Fig. 6E), consistent with our observation that there was no further increase in BCMA expression (Fig 6D). We were not able to test the effect of compound combinations on HDP101 sensitivity in AMO1 and RPMI-8226 cell lines, since these lines are not sensitive to HDP101.

**Fig 6:**
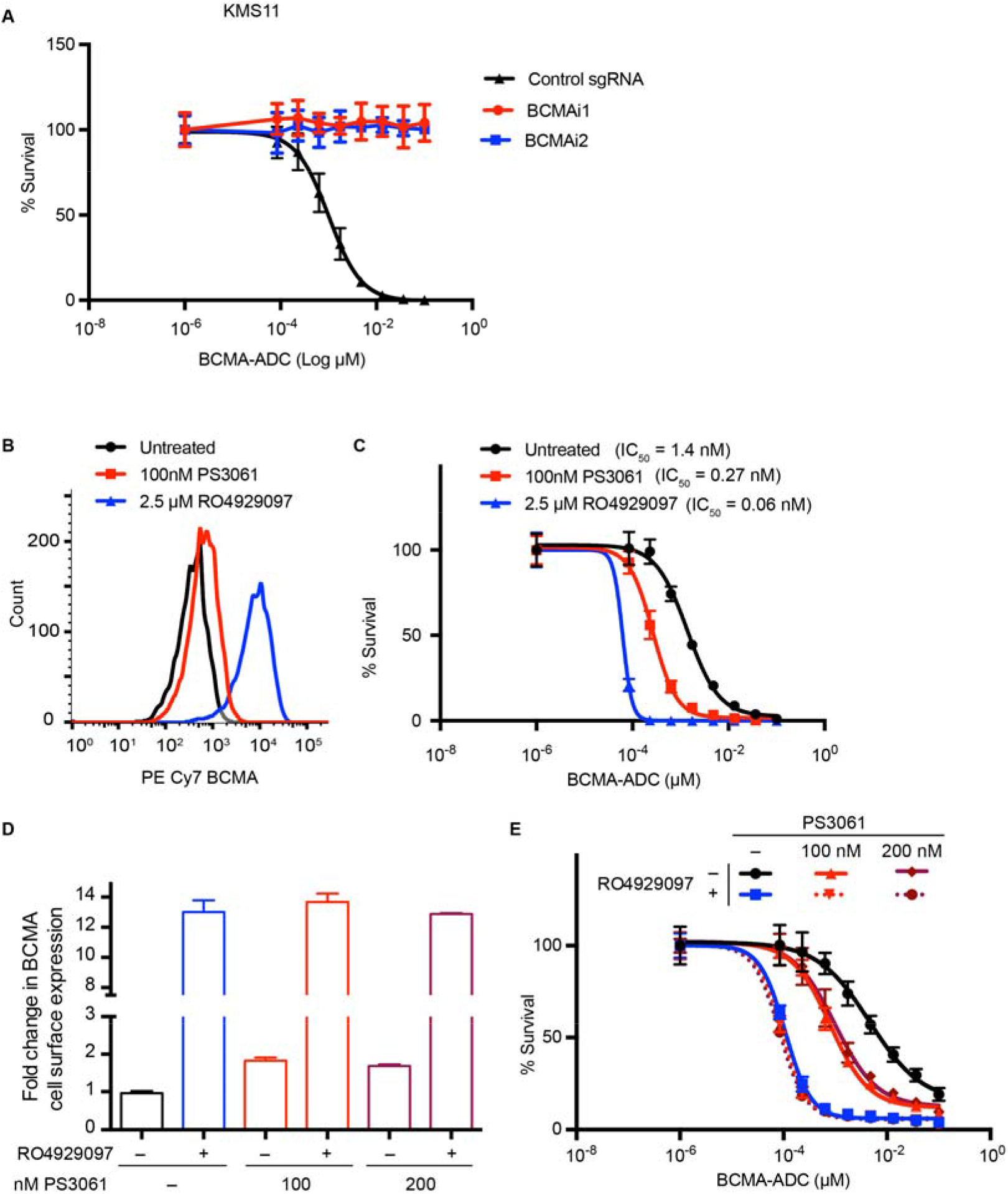
Increased efficacy of BCMA-ADC when combined with Sec61 inhibitor. **A**, BCMA-ADC cytotoxicity dose-response assay of CRISPRi KMS11 cells expressing two independent sgRNA targeted towards BCMA and a non-targeting control sgRNA. **B,** KMS11 cells treated with indicated concentrations of PS3061 and RO4929097 were stained for BCMA and analyzed by flow cytometry. Histograms indicate the distribution of BCMA. Data is a representation of two biological replicates. **C,** BCMA-ADC cytotoxicity dose-response assay was performed in combination with indicated concentrations of drugs. Data points are means of two biological replicates; error bars denote SD. **D,** KMS11 cells treated with indicated concentrations of PS3061 and 2.5 μM RO4929097 were stained for BCMA and analyzed by flow cytometry. Data is a representation of two biological replicates; error bars denote SD. **E,** BCMA-ADC cytotoxicity dose-response assay was performed in combination with indicated concentrations of drugs. Data points are means of two biological replicates; error bars denote SD.

Taken together, these findings demonstrate that our CRISPR screen was able to identify druggable targets that enhance the efficacy of immunotherapy agents.

### BCMA CAR-T cell co-culture screen reveals myeloma cell-intrinsic proteins that affect CAR-T efficacy

A systematic understanding of mechanisms causing resistance to CAR-T cell therapy will enable us to design potential combination therapies pre-empting resistance. Here, we used our CRISPR screening platform to identify genes or pathways that control sensitivity and resistance of MM cells to BCMA-targeted CAR-T cells.

First, we generated BCMA CAR-T cells by transducing CD8+ T cells with a lentiviral vector encoding a second generation CAR incorporating an anti-BCMA single chain variable fragment, 4-1BB costimulatory domain and CD3-zeta signaling domain (see Methods). We validated the activation of BCMA CAR-T cells in the presence of AMO1 cells (Fig. 7A), and their cytotoxicity against AMO1 cells (Fig. 7B). Knockdown of BCMA in AMO1 cells using two independent sgRNAs (Supplemental Fig. 5) reduced both activation and cytotoxicity of BCMA-targeted CAR-T cells (Fig. 7A-B). This finding indicated that the efficacy of the CAR-T cells depends on the cell-surface levels of BCMA in MM cells (Fig. 7A-B).

**Fig 7:**
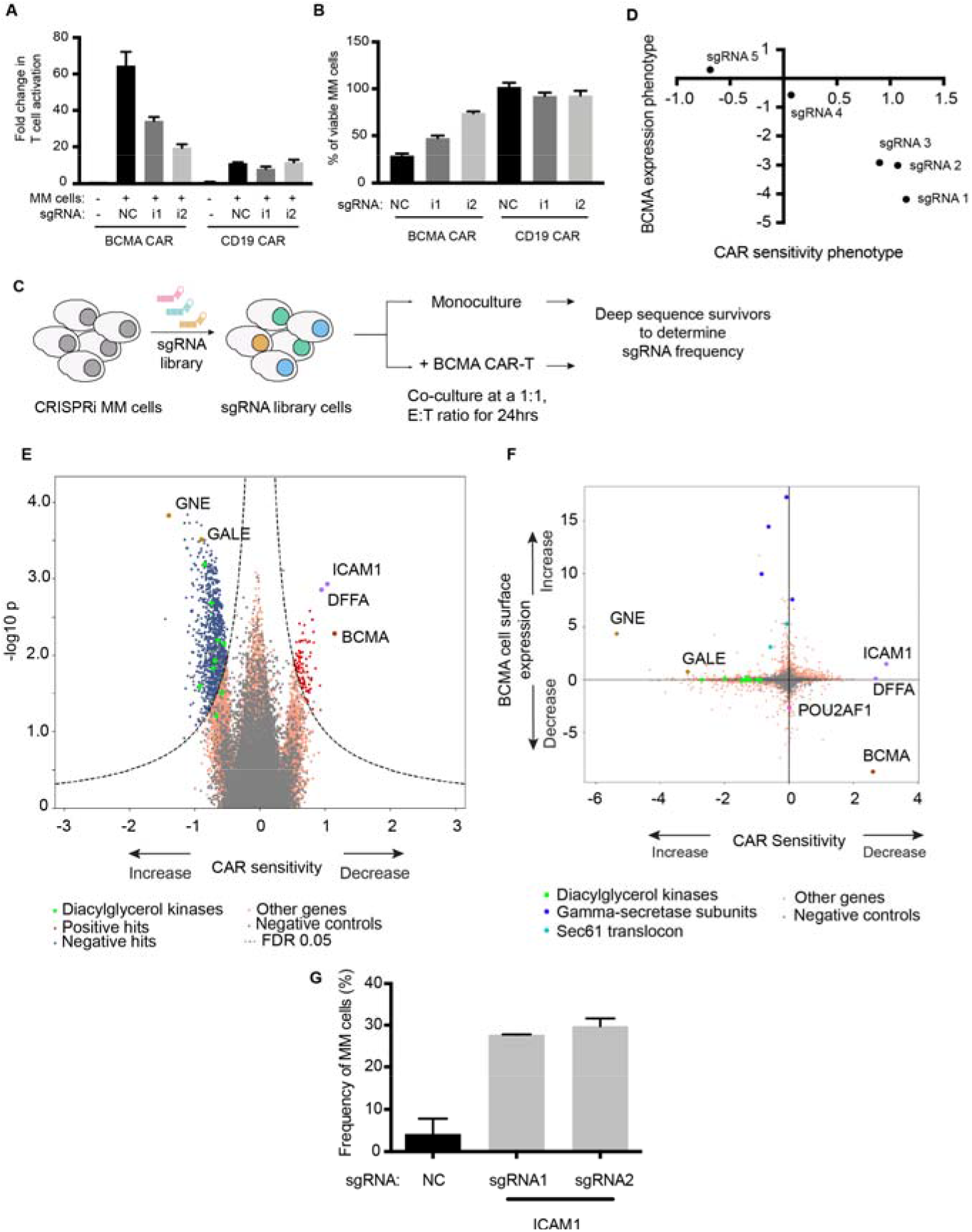
CRISPRi-CAR-T screen identifies genes modulating response to BCMA CAR-T cells. **A and B,** CRISPRi AMO1 cells expressing BFP-sgRNA targeted towards BCMA or nontargeting control sgRNA were co-cultured at a 1:1 ratio with BCMA- or CD19- GFP CAR-T cells. Fold changes in T cell activation was determined by analyzing for cell surface expression of CD69 on T cells normalized to CD69 expression on resting T cells. Myeloma cell viability was determined by propidium iodide staining of BFP-positive myeloma cells. **C,** Schematic representation of the CRISPRi-CAR-T cell co-culture screen. **D,** Comparison of BCMA expression phenotype to sensitivity towards BCMA-CAR indicating different sgRNAs targeted towards BCMA. **E,** Volcano plot indicating CAR-T sensitivity phenotypes and statistical significance of knockdown of human genes (orange dots) and quasi-genes from negative control sgRNAs (grey dots) in multiple myeloma cells. Hits genes corresponding to functional categories are color-coded as labeled in the panel. **F,** Comparison of BCMA expression phenotype from the CRISPRi primary screen (Fig. 1) to sensitivity towards BCMA-CAR. Hits genes corresponding to functional categories are color-coded as labeled in the panel. **G,** CRISPRi AMO1 cells expressing two independent sgRNAs targeting *ICAM1* or non-targeting control sgRNA were cocultured overnight at a 1:1 ratio with BCMA-CAR-T cells. Frequency of viable myeloma cells was determined by flow cytometry.

While the expression level of the target antigen is a major determinant of response to CAR-T cells, there are likely also antigen-independent determinants. To systematically identify such determinants, we conducted a CRISPRi screen to identify genes or pathways in MM cells determining response to BCMA CAR-T cells (Fig. 7C). AMO1 cells expressing the CRISPRi machinery and an sgRNA library targeting 12,838 genes (including kinases, phosphatases, cancer drug targets, apoptosis genes, mitochondrial genes and transcription factors) were grown as mono-culture or co-cultured with BCMA CAR-T cells. 24 hours later, the surviving cells were harvested and processed for next-generation sequencing.

This CRISPRi screen identified a substantial number of genes regulating sensitivity to CAR-T cells (Fig. 7E and Supplemental Table 6). Comparing BCMA cell-surface levels to CAR-T sensitivity for different sgRNAs targeting BCMA, we observed that surface levels correlated with sensitivity to BCMA CAR-T cells (Fig. 7D). For example, knockdown of subunits of the gamma secretase complex and genes involved in sialic acid biosynthesis pathways *GALE* and *GNE*, which change BCMA cell surface expression, also concordantly affected sensitivity to BCMA CAR-T cells (Fig. 7E-F).

Furthermore, the screen identified a different category of genes, knockdown of which affected sensitivity of AMO1 cells to BCMA CAR-T cells without affecting cell-surface levels of BCMA. These genes included intercellular adhesion molecule 1 (*ICAM1*), which functions in T cell activation^38^, and DNA fragmentation factor subunit alpha (*DFFA*), which is required for caspase activation to trigger apoptosis^39^. Knockdown of *ICAM1* or *DFFA* resulted in decreased CAR-T cell sensitivity. Conversely, knockdown of genes belonging to the family of diacyl glycerol kinases (DGK) caused increased sensitivity to BCMA CAR-T cells (Fig. 7F). Using two individual sgRNAs targeting *ICAM1*, we were able to validate that knockdown of ICAM1 results in reduced sensitivity to BCMA CAR-induced cytotoxicity (Fig. 7G).

This screen thus identified potential pathways that can regulate sensitivity to CAR-T cells either through changing cell surface expression of the antigen, or through an antigen-independent mechanism.

## Discussion

Our study establishes that CRISPR-based screens enable a systematic and scalable approach to identifying mechanisms of antigen expression. Uncovering pathways that regulate BCMA transcription, translation and trafficking to the cell surface enables us to propose effective treatment strategies in patients with low basal expression of BCMA or following relapse due to antigen-loss. Clinical trials are currently ongoing to investigate the combination of gamma-secretase inhibitor with BCMA CAR-T cell therapy (NCT03502577). However, it is likely that not all patients would respond similarly to gamma secretase inhibition, or resistance may also develop to this combination strategy. Therefore, identifying alternative ways to modulate BCMA, as we discover here, retains high clinical relevance. Our results also support the concept that strategies to upregulate antigen expression on cancer cells can enhance efficacy of antigen-targeted immunotherapy agents.

Our findings also raise several mechanistic questions, which will be subject of future studies. In particular, it is not clear how SEC61 inhibition results in increased cell-surface levels of BCMA. SEC61 knockdown was previously shown to upregulate a subset of proteins^40^. This effect on BCMA could be due to direct effects on the interaction of BCMA with the SEC61 complex, or due to indirect effects mediated by changes in other SEC61 clients.

Furthermore, our results indicate HDAC7 as a potential combination target for BCMA-targeted immunotherapy. This highlights the importance of inhibiting specific HDACs, and provides a potential therapeutic use case for an HDAC7-specific inhibitor to enhance BCMA-targeted immunotherapy.

Using a complementary strategy, our CRISPRi-CAR-T coculture screen uncovered both antigen-dependent and independent mechanisms in MM cells that control the response to BCMA-targeted CAR-T cells. Recent screens for pathways controlling cytotoxicity of CAR-T cells in leukemia and lymphoma cells showed death receptor signaling to play an important role^12^, whereas our screens did not find the same pathway. This could be due to differences in methodology, CAR target antigens (BCMA versus CD19), or the biology of the different cancer types. Our screen showed that knockdown of *DGK* family of genes resulted in increased sensitivity to CAR-T cells. Previous groups have shown that inhibition of DGK-α activity in T cells using pharmacological inhibitors induced T cell activation, thus improving its cytotoxic activity on cancer cells^41,42^. Our study indicates that knockdown of *DGK* kinases in MM cells increases sensitivity to T cells. This indicates that inhibitors against this family of kinases may be beneficial in CAR-T cell therapy of MM through a dual mechanism, acting by both increasing T cell activity but also sensitizing MM cells to CAR-T cells

Our study also identified genes in the sialic acid biosynthesis pathways, *GALE* and *GNE*, knockdown of which sensitized MM cells to CAR-T cells. Although our BCMA-expression screen identified these genes to upregulate BCMA, the effect size of sensitizing the cells to CAR-T cells was much higher than the change in BCMA expression, indicating that there could be additional mechanisms by which this pathway modulates response to CAR-T cells. Sialic acid blockade in cancer cells has been shown to create an immune-permissive tumor microenvironment increasing CD8+ T cell activity^43^. Therefore, inhibition of GALE and/or GNE could modulate the tumor microenvironment and increase sensitivity to CAR-T cells in an antigen-independent mechanism.

In summary, we present two complementary screening approaches to identify potential combinatorial treatments that can enhance the efficacy of immunotherapy through antigen-dependent and -independent mechanisms. These strategies should be readily adaptable to a broad range of cancer cell types, antigens, and immunotherapy modalities.

## Acknowledgements

This work was supported by a Postdoctoral Fellowship from the UCSF Program for Breakthrough Biomedical Research (to P.R.), K99/R00 CA181494 (to M.K.), a Stand Up to Cancer Innovative Research Grant (to M.K.), the UCSF Stephen and Nancy Multiple Myeloma Translational Initiative (to M.K., K.T.R, A.P.W), UCSF Breast Cancer Research Funds (to J.T.), R01 CA226851 (to A.P.W.).

We thank James Nunez, Marco Jost, Christina Liem and Jonathan Weissman for sharing their unpublished CRISPRa construct; Diego Acosta-Alvear for sharing mCherry-Luciferase RPMI8226 cell line; Axel Hyrenius-Wittsten and Joe Hiatt for inputs on T cell culturing; Eric Chow and Derek Bogdanoff (UCSF Center for Advanced Technology) for support with nextgeneration sequencing; Sarah Elmes and Jane Gordon (UCSF Laboratory for Cell Analysis) for support with FACS; Stratton Georgoulis, Molly Bassette and Logan Hille for contributing to preliminary studies. For primary patient samples, we thank Nina Shah, Sandy W. Wong, Tom G. Martin III, Jeffrey L. Wolf and the Grand Multiple Myeloma Translational Initiative. We thank all the co-authors and members of the Kampmann lab for discussion and feedback on the manuscript.

## Authorship contributions

P.R., A.P.W, M.K designed the study; P.R., M.S., J.T.L., M.C., K.T.R. developed methodology; P.R., A.B.A., M.S., J.T.L., M.C. performed the experiments; P.R., A.B.A., R.T., J.T., A.P.W., M.K. analyzed the data; J.T.L., P.C., T.H., N.S., S.W.W., T.G.M., J.L.W., K.T.R., A.P., J.T. provided material and technical support; P.R., M.K. wrote the manuscript; and all co-authors approved final version of the manuscript.

## Data Sharing Statement

For original data not included in the manuscript and supplemental tables, please contact.

## References

1. Cho SF, Anderson KC, Tai YT. Targeting B Cell Maturation Antigen (BCMA) in Multiple Myeloma: Potential Uses of BCMA-Based Immunotherapy. Front Immunol. 2018;9:1821.

2. Raje N, Berdeja J, Lin Y, et al. Anti-BCMA CAR T-Cell Therapy bb2121 in Relapsed or Refractory Multiple Myeloma. N Engl J Med. 2019;380(18):1726–1737.

3. Susanibar Adaniya SP, Cohen AD, Garfall AL. Chimeric antigen receptor T cell immunotherapy for multiple myeloma: A review of current data and potential clinical applications. Am J Hematol. 2019;94(S1):S28–S33.

4. Brudno JN, Maric I, Hartman SD, et al. T Cells Genetically Modified to Express an Anti-B-Cell Maturation Antigen Chimeric Antigen Receptor Cause Remissions of Poor-Prognosis Relapsed Multiple Myeloma. J Clin Oncol. 2018;36(22):2267–2280.

5. Cohen AD, Garfall AL, Stadtmauer EA, et al. B cell maturation antigen-specific CAR T cells are clinically active in multiple myeloma. J Clin Invest. 2019;129(6):2210–2221.

6. Majzner RG, Mackall CL. Tumor Antigen Escape from CAR T-cell Therapy. Cancer Discov. 2018;8(10):1219–1226.

7. Iorgulescu JB, Braun D, Oliveira G, Keskin DB, Wu CJ. Acquired mechanisms of immune escape in cancer following immunotherapy. Genome Med. 2018;10(1):87.

8. Nijhof IS, Casneuf T, van Velzen J, et al. CD38 expression and complement inhibitors affect response and resistance to daratumumab therapy in myeloma. Blood. 2016;128(7):959–970.

9. Pont MJ, Hill T, Cole GO, et al. gamma-secretase inhibition increases efficacy of BCMA-specific chimeric antigen receptor T cells in multiple myeloma. Blood. 2019.

10. Han P, Dai Q, Fan L, et al. Genome-Wide CRISPR Screening Identifies JAK1 Deficiency as a Mechanism of T-Cell Resistance. Front Immunol. 2019;10:251.

11. Tsui CK, Barfield RM, Fischer CR, et al. CRISPR-Cas9 screens identify regulators of antibody-drug conjugate toxicity. Nat Chem Biol. 2019;15(10):949–958.

12. Dufva O, Koski J, Maliniemi P, et al. Integrated drug profiling and CRISPR screening identify essential pathways for CAR T-cell cytotoxicity. Blood. 2020;135(9):597–609.

13. Burr ML, Sparbier CE, Chan YC, et al. CMTM6 maintains the expression of PD-L1 and regulates anti-tumour immunity. Nature. 2017;549(7670):101–105.

14. Dong MB, Wang G, Chow RD, et al. Systematic Immunotherapy Target Discovery Using Genome-Scale In Vivo CRISPR Screens in CD8 T Cells. Cell. 2019;178(5):1189–1204 e1123.

15. Gilbert LA, Horlbeck MA, Adamson B, et al. Genome-Scale CRISPR-Mediated Control of Gene Repression and Activation. Cell. 2014;159(3):647–661.

16. Horlbeck MA, Gilbert LA, Villalta JE, et al. Compact and highly active next-generation libraries for CRISPR-mediated gene repression and activation. Elife. 2016;5.

17. Kampmann M, Bassik MC, Weissman JS. Functional genomics platform for pooled screening and generation of mammalian genetic interaction maps. Nat Protoc. 2014;9(8):1825–1847.

18. Tian R, Gachechiladze MA, Ludwig CH, et al. CRISPR Interference-Based Platform for Multimodal Genetic Screens in Human iPSC-Derived Neurons. Neuron. 2019.

19. Eisen MB, Spellman PT, Brown PO, Botstein D. Cluster analysis and display of genomewide expression patterns. Proc Natl Acad Sci U S A. 1998;95(25):14863–14868.

20. Saldanha AJ. Java Treeview--extensible visualization of microarray data. Bioinformatics. 2004;20(17):3246–3248.

21. Figueroa-Vazquez V. KJ, Breunig C., Baumann, A., Giesen N., Pálfi A., Müller C., Lutz C., Hechler T., Kulke M., Mueller-Tidow C., Goldschmidt H., Pahl A., Raab MS. HDP-101, a novel anti-BCMA antibody-drug conjugate, specifically kills proliferating and resting multiple myeloma cells. Submitted.

22. Pahl A, Ko J, Breunig C, et al. HDP-101: Preclinical evaluation of a novel anti-BCMA antibody drug conjugates in multiple myeloma. Journal of Clinical Oncology. 2018;36(15_suppl):e14527–e14527.

23. Imai C, Mihara K, Andreansky M, et al. Chimeric receptors with 4-1BB signaling capacity provoke potent cytotoxicity against acute lymphoblastic leukemia. Leukemia. 2004;18(4):676–684.

24. Jost M, Chen Y, Gilbert LA, et al. Combined CRISPRi/a-Based Chemical Genetic Screens Reveal that Rigosertib Is a Microtubule-Destabilizing Agent. Mol Cell. 2017;68(1):210–223 e216.

25. Zhao C, Inoue J, Imoto I, et al. POU2AF1, an amplification target at 11q23, promotes growth of multiple myeloma cells by directly regulating expression of a B-cell maturation factor, TNFRSF17. Oncogene. 2008;27(1):63–75.

26. Laurent SA, Hoffmann FS, Kuhn PH, et al. gamma-Secretase directly sheds the survival receptor BCMA from plasma cells. Nat Commun. 2015;6:7333.

27. Luistro L, He W, Smith M, et al. Preclinical profile of a potent gamma-secretase inhibitor targeting notch signaling with in vivo efficacy and pharmacodynamic properties. Cancer Res. 2009;69(19):7672–7680.

28. Ding ZM, Liu YG. [Effects of phosphoramidothiolate pesticides on rat erythrocyte membrane acetylcholinesterase]. Zhonghua Yu Fang Yi Xue Za Zhi. 1988;22(2):82–84.

29. Lobera M, Madauss KP, Pohlhaus DT, et al. Selective class IIa histone deacetylase inhibition via a nonchelating zinc-binding group. Nat Chem Biol. 2013;9(5):319–325.

30. Scuto A, Kirschbaum M, Kowolik C, et al. The novel histone deacetylase inhibitor, LBH589, induces expression of DNA damage response genes and apoptosis in Ph-acute lymphoblastic leukemia cells. Blood. 2008;111(10):5093–5100.

31. Santo L, Hideshima T, Kung AL, et al. Preclinical activity, pharmacodynamic, and pharmacokinetic properties of a selective HDAC6 inhibitor, ACY-1215, in combination with bortezomib in multiple myeloma. Blood. 2012;119(11):2579–2589.

32. Choi SY, Kee HJ, Jin L, et al. Inhibition of class IIa histone deacetylase activity by gallic acid, sulforaphane, TMP269, and panobinostat. Biomed Pharmacother. 2018;101:145–154.

33. Mackinnon AL, Paavilainen VO, Sharma A, Hegde RS, Taunton J. An allosteric Sec61 inhibitor traps nascent transmembrane helices at the lateral gate. Elife. 2014;3:e01483.

34. Garrison JL, Kunkel EJ, Hegde RS, Taunton J. A substrate-specific inhibitor of protein translocation into the endoplasmic reticulum. Nature. 2005;436(7048):285–289.

35. Shah PS, Link N, Jang GM, et al. Comparative Flavivirus-Host Protein Interaction Mapping Reveals Mechanisms of Dengue and Zika Virus Pathogenesis. Cell. 2018;175(7):1931–1945 e1918.

36. Collins DM, Bossenmaier B, Kollmorgen G, Niederfellner G. Acquired Resistance to Antibody-Drug Conjugates. Cancers (Basel). 2019;11(3).

37. Pahl A, Lutz C, Hechler T. Amanitins and their development as a payload for antibodydrug conjugates. Drug Discov Today Technol. 2018;30:85–89.

38. Wingren AG, Parra E, Varga M, et al. T cell activation pathways: B7, LFA-3, and ICAM-1 shape unique T cell profiles. Crit Rev Immunol. 1995;15(3-4):235–253.

39. Gu J, Dong RP, Zhang C, McLaughlin DF, Wu MX, Schlossman SF. Functional interaction of DFF35 and DFF45 with caspase-activated DNA fragmentation nuclease DFF40. J Biol Chem. 1999;274(30):20759–20762.

40. Nguyen D, Stutz R, Schorr S, et al. Proteomics reveals signal peptide features determining the client specificity in human TRAP-dependent ER protein import. Nat Commun. 2018;9(1):3765.

41. Noessner E. DGK-alpha: A Checkpoint in Cancer-Mediated Immuno-Inhibition and Target for Immunotherapy. Front Cell Dev Biol. 2017;5:16.

42. Riese MJ, Moon EK, Johnson BD, Albelda SM. Diacylglycerol Kinases (DGKs): Novel Targets for Improving T Cell Activity in Cancer. Front Cell Dev Biol. 2016;4:108.

43. Bull C, Boltje TJ, Balneger N, et al. Sialic Acid Blockade Suppresses Tumor Growth by Enhancing T-cell-Mediated Tumor Immunity. Cancer Res. 2018;78(13):3574–3588.

