## Supplemental Material for "CRISPR-based screens uncover determinants of immunotherapy response in Multiple Myeloma"

### Supplemental Materials

**Supplemental Methods**

**Legends for Supplemental Tables 1-6**

**Supplemental Tables 7,8**

**Supplemental Figures 1-5**

**Supplemental references**

**Supplemental Methods:**

### Cell culture

Multiple myeloma cell lines AMO1 (gift from Christoph Driessen, Kantonsspital St. Gallen, Switzerland) , KMS11 (JCRB), KMS12-PE (JCRB), OPM2 (DSMZ) and RPMI8226-luciferase (gift from Diego Acosta-Alvear, UCSB) [1] were grown in RPMI1640 complete growth media containing 10% fetal bovine serum (# 97068-085, Lot# 076B16, Seradigm), 1% Penicillin/Streptomycin (#15140122, Life tech) and 2 mM L-Glutamine (#25030081, Life tech). Cell lines were seeded at 0.2-0.4*$10^6$ cells/ml and[2,3] passaged when they reached 1*$10^6$ cells/ml cell density. All cell lines were validated by STR profiling service provided by Genetica DNA Laboratories.

### Drug treatment of cells and downstream assays

For drug treatment, MM cells were seeded at a concentration of 0.2*$10^6$ cells/ml and treated with indicated concentration of TMP269 (#S7324; Selleckchem), Panobinostat (#S1030; Selleckchem), Ricolinostat (#S8001; Selleckchem), RO4929097 (#S1575; Selleckchem) for 48 hrs and with the Sec61 inhibitors, CT8 [4] and PS3061[5] for 24 hrs. Following drug treatment, cells were harvested and analyzed by flow cytometry for cell-surface expression, immunoblotting for total protein levels and qPCR for transcript levels of BCMA and CD38. The assays are detailed in supplemental materials and methods.

### Multiple myeloma patient culture and drug treatments

De-identified primary MM bone marrow aspirates were obtained from the UCSF hematological malignancy tissue bank in compliance with the UCSF IRB protocols. Bone marrow mononuclear cells were isolated by density gradient centrifugation using Histopaque-1077 (#10771, Sigma-Aldrich). Briefly, 3 ml of bone marrow aspirate was diluted with D-PBS without Calcium and Magnesium to a final volume of 8 ml and carefully laid over 3 ml of Histopaque-1077 and centrifuged at 400 g for 30 mins at room temperature. The layer containing plasma cells was carefully isolated, washed with PBS and resuspended in RPMI complete growth media to a concentration of 0.5-1*$10^6$ cells/ml. For drug treatment, cells were treated in triplicates for 24 hrs with indicated concentrations of drugs or DMSO. Following drug treatments, cells were stained using PE/Cy7-BCMA (19F2), FITC-CD38 (HIT2), APC-R700-CD138 (MI15) (#566050, BD Biosciences) antibodies and propidium iodide or their corresponding isotype control antibodies and analyzed by flow cytometry using Bio-Rad ZE5 cell analyzer.

### sgRNA libraries

The genome-wide CRISPRi and CRISPRa v2.0 libraries subdivided into seven sublibraries containing five sgRNAs per gene were described previously[6]. For pooled screening, sgRNA libraries were packaged into lentivirus for transduction into MM cell lines as follows. HEK293T cells were transfected with the sgRNA sublibrary [6] and third generation packaging mix [7] using Mirus TransIT-Lenti reagent (#MIR6600, Mirus Bio). 48 hrs later, the viral supernatant was harvested, filtered through a 0.45 μm filter and precipitated using Alstem Lenti precipitating solution (#VC100, Alstem, Inc.).

### CRISPR cell line generation

To establish cell lines expressing functional CRISPRi, the MM cell lines were lentivirally transduced with pMH0006 [8] and polyclonal cell lines were established by fluorescence-activated cell sorting (FACS) for BFP-positive cells. CRISPRi activity was validated using sgRNA targeted towards a cell surface marker, *CD38* (Supplementary Table S6). Briefly, CRISPRi MM cells expressing CD38 sgRNA or non-targeting control sgRNA were stained using PE/Cy7-CD38 (HIT2) antibody (#303515, Biolegend) and analyzed for changes in levels of CD38 by flow cytometry using BD FACSCelesta™ (BD Biosciences).

For establishing CRISPRa cell line, AMO1 cells were lentivirally transduced with pMJ467-pHR-dCas9-HA-XTEN-VPR-2XP2A-mCherry (gift from James Nunez, Marco Jost and Jonathan Weissman, sequence available at (kampmannlab.ucsf.edu/resources). This plasmid is an optimized version of JKNp44 [9], in which a universal chromatin opening element was included upstream of the SFFV promoter, the linker between dCas9 and the VPR domain was replaced by an XTEN linker, and BFP was replaced by mCherry. Transduced cells were flow sorted based on mCherry fluorescence to generate a polyclonal cell line. CRISPRa activity was determined as described for CRISPRi using sgRNA targeted towards cell surface marker, *CXCR4* (Supplementary Table S6).

**Staining for flow cytometry**

Drug treated cells were stained with PE/Cy7-BCMA (19F2), FITC-CD38 (HIT2) and propidium iodide (#P3566, Invitrogen) or corresponding isotype control antibodies (#400253; #400107, Biolegend) and analyzed by flow cytometry using BD FACS Celesta™ (BD Biosciences). Flow cytometry data was analyzed using FlowJo v10.4 (FlowJo), raw median fluorescence intensity (MFI) values of BCMA and CD38 stained cells were normalized to isotype control samples and data plotted as fold change in MFI relative to untreated cells using Prism V7 (GraphPad Software).

**Western blotting**

Total cell lysates were collected by resuspending cells in RIPA buffer (#89900, Thermo Fisher Scientific) supplemented with 1X protease inhibitor cocktail (#11836170001, Roche). Protein concentration was determined using BCA assay (#23225, Thermo Fisher Scientific) and protein extracts were prepared in SDS sample buffer. Equal concentrations of the cell lysates were loaded onto a NuPage 4-12% Bis-tris gels (#NP0336BOX, Thermo Fisher Scientific) and transferred onto a 0.45 μm pore nitrocellulose membrane using the Trans-Blot Turbo Transfer System (#1704270, Bio-Rad Laboratories, Inc.). The primary antibodies used in this study include: rabbit anti-BCMA (E6D7B) (#88183, Cell Signaling Technology), 1:2000; mouse anti-GAPDH (0411) (#sc-47724, Santa Cruz Biotechnology), 1:5000. Blots were incubated with Li-Cor secondary antibodies and imaged using Odyssey Fc imaging system (#2800, Li-Cor). Digital images were processed and analyzed using Li-Cor Image Studio software.

**Quantitative-PCR (qPCR)**

Drug treated cells or CRISPRi cells expressing sgRNA were harvested by centrifugation, and RNA extracted using the Quick-RNA miniprep kit (#R1054, Zymo Research). 1 μg of total RNA was reverse transcribed using Superscript III First-Strand Synthesis system

(#18080051, Invitrogen). qPCR was performed using SensiFast SYBR Lo-ROX 2X qPCR master mix (#BIO-94005, Bioline) following manufacturers guidelines. Each qPCR reaction was set up in triplicates and run on Quantstudio 6 Flex (Applied Biosystems) following manufacturers protocol. Expression fold changes were calculated using the delta-delta Ct method and normalized to an internal control (beta-actin). The qPCR primers used are listed in Supplementary Table S6.

**ELISA assay**

To determine soluble BCMA (sBCMA) levels in the cell culture supernatant, solid phase sandwich ELISA using polyclonal goat antibodies was used (#DY193, Human BCMA/TNFRSF17 Duoset ELISA, R&D Systems). Cell culture supernatant was harvested from cells treated with indicated concentration of drugs and concentration of sBCMA was determined following the manufacturer's instructions.

**Isolation of CD8+ T cells**

De-identified donor blood after apheresis was obtained from Blood Centers of the Pacific (San Francisco, CA) as approved by UCSF IRB policies. CD8+ T cells were isolated from donor blood using EasySep ™ Human CD8+ T cell isolation kit (#19053, Stemcell Technologies) following the manufacturers protocol. Purified CD8+ T cells were cryopreserved in RPMI1640 media supplemented with 20% human AB serum (#HP1022, Valley Medical) and 10% DMSO (#D2650, Sigma) solution. For experiments, T cells were cultured in Ex-Vivo 15 media (#04-418Q, Lonza) supplemented with 5% human AB serum, 10mM neutralized N-acetyl L-Cysteine (#A9165, Sigma Aldrich), 1X beta-mercaptoethanol (#21985-023, Thermo Fisher Scientific) and 50 units/ml of recombinant human IL-2 (#200-02, Peprotech).

**Generation of CAR construct and CAR-T cells**

To generate the BCMA-targeted CAR, the nucleotide sequence for the BCMA-50 scFv (Patent: https://patents.google.com/patent/US20170183418) and a N-terminal CD8a signal peptide plus myc-tag was synthesized. Using Gibson cloning, the CAR was assembled by fusing the BCMA-50 scFv to the CD8a hinge and transmembrane domain, 4-1BB costimulatory domain, the CD3-zeta chain, and a C-terminal EGFP. The CAR was then inserted into the second generation lentiviral vector pHR-SIN. The CD19 CAR has been previously described [4].

For CAR-T cell generation, lentivirus was produced in HEK293T cells by transfecting pHR'SIN:CAR-Transgene vector and lentiviral packaging plasmids, pMD2.G and pCMVdR8.9 using Mirus TransIT-Lenti reagent. Viral supernatant was harvested 48 hr after transfection and precipitated using Alstem lentivirus precipitating solution. Primary CD8+ T cells were thawed and 24 hrs later stimulated with Dynabeads Human T-Expander CD3/CD28 beads (#11141D, ThermoFisher Scientific) at a cell:bead ratio of 1:2. The following day, primary T cells were exposed to the precipitated virus for 24 hrs. On day 5 post stimulation, Dynabeads were removed and GFP CAR-T cells were enriched by FACS sorting. The sorted CAR-T cells were maintained in culture until day 10-12 when they are used for downstream assays.

**Legends for Supplemental Tables 1-6**

**Supplemental Table 1** – Results from the genome-wide CRISPRi screen for BCMA cell-surface levels. Phenotypes and p values were calculated using the MAGeCK-iNC pipeline [17]. Columns are: A. TSS - Targeted transcription start site, B: Epsilon - knockdown phenotype for screen where negative values indicate downregulation of BCMA and positive values indicate upregulation of BCMA, C: p Value, D: product (epsilon x –log10(p value)), E: Gene name

**Supplemental Table 2** - Results from the genome-wide CRISPRa screen for BCMA cell-surface levels. Phenotypes and p values were calculated using the MAGeCK-iNC pipeline [17]. Columns are: A. TSS - Targeted transcription start site, B: Epsilon - knockdown phenotype for screen where negative values indicate downregulation of BCMA and positive values indicate upregulation of BCMA, C: p Value, D: product (epsilon x –log10(p value)), E: Gene name

**Supplemental Table 3** – Results from the CRISPRi validation screens for individual sgRNAs. Phenotypes were calculated using the CRISPR bioinformatics pipeline as described previously [17]. Columns are: A. Gene name, B. sgRNA targeting the gene, C. sgRNA protospacer sequence, D-M. Knockdown phenotype score for changes in BCMA or CD38 in a panel of CRISPRi myeloma cell lines.

**Supplemental Table 4** – Results from the CRISPRi validation screen for genes. Phenotypes were calculated using the CRISPR bioinformatics pipeline as described previously [17]. Sheet1. CRISPRi validation phenotype for both BCMA and CD38 Columns are: A. Gene name, B-K. Knockdown phenotype score for changes in BCMA or CD38 in a panel of CRISPRi myeloma cell lines. Sheet 2. Average BCMA phenotype and ranking of positive hit genes in the CRISPRi validation screen. Columns are: A. Gene name, B-F. Knockdown phenotype score for changes in BCMA in a panel of CRISPRi myeloma cell lines, G. Average BCMA phenotype for MM cell lines, H. Ranking. Sheet 3. Average BCMA phenotype and ranking of negative hit genes in the CRISPRi validation screen. Columns are: A. Gene name, B-F. Knockdown phenotype score for changes in BCMA in a panel of CRISPRi myeloma cell lines, G. Average BCMA phenotype for MM cell lines, H. Ranking.

**Supplemental Table 5** – Multiple myeloma patient information. Columns are: A. Patient number B. Sex C. Age D. Disease status during sample collection E. Fluorescence in situ hybridization (FISH) analysis of patient bone marrow aspirate during sample collection

**Supplemental Table 6** – Results from the CRISPRi CAR-T cell coculture survival screen. Phenotypes and p values were calculated using the MAGeCK-iNC pipeline [17]. Columns are: A. TSS - Targeted transcription start site, B: Epsilon - knockdown phenotype for screen where negative values indicate downregulation of BCMA and positive values indicate upregulation of BCMA, C: p Value, D: product (epsilon x –log10(p value)), E: Gene name

| sgRNA short name | Gene | protospacer sequence | Mode of perturbation |
|---|---|---|---|
| BCMA-sgRNA1 | TNFRSF17 | GAATAACGCTGACATGTTAG | CRISPRi |
| BCMA-sgRNA2 | TNFRSF17 | GCTGCTTCGTGGGTTCTTAC | CRISPRi |
| CD38 | CD38 | GGGCTGGGCGAAGATGAGGC | CRISPRi |
| CXCR4 | CXCR4 | GCAGACGCGAGGAAGGAGGGCGC | CRISPRa |

**Supplemental Table 7** – List of sgRNA sequences. Columns are: A. sgRNA name, B. Gene name, C. protospacer sequence, D. Mode of perturbation.

| Gene | Orientation | Sequence (5' to 3') |
|---|---|---|
| BCMA | Forward | CTTGCATACCTTGTCAACTTCG |
| | Reverse | TTAAGCTCAGTCCCAAACAGG |
| CD38 | Forward | GTATTGGTTGAAAGGAGTGCTG |
| | Reverse | CACGCTAACTGGTTGAATCTCT |
| Beta-Actin | Forward | ACCTTCTACAATGAGCTGCG |
| | Reverse | CCTGGATAGCAACGTACATGG |
| GAPDH | Forward | ATGCCTCCTGCACCACCAAC |
| | Reverse | GGGGCCATCCACAGTCTTCT |

**Supplemental Table 8** – Oligonucleotides for qPCR analysis. Columns are: A. Gene name, B. Orientation of the sequence, C. primer sequence (5’ -> 3’).

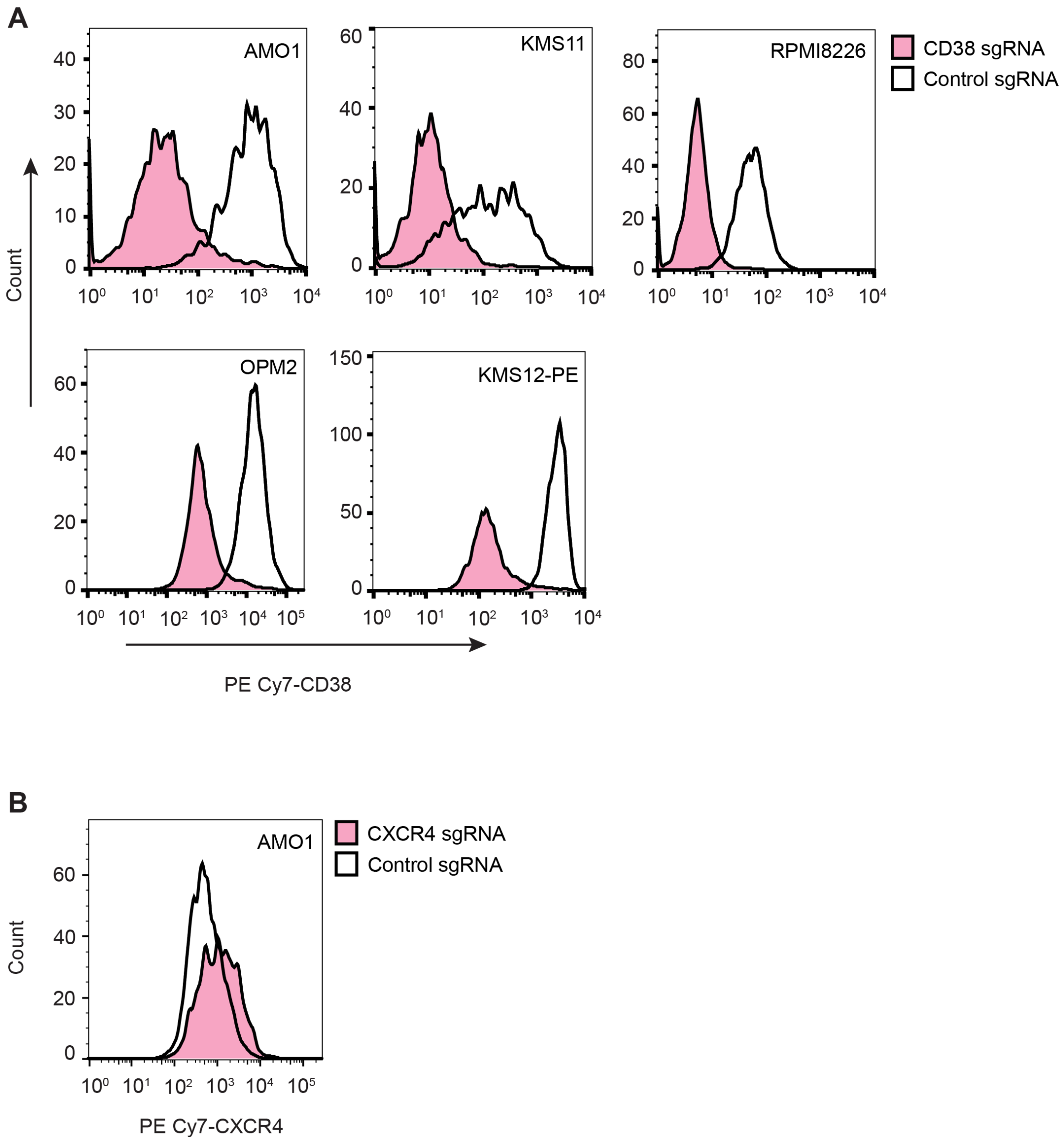


**Supplemental Fig 1:**

**Validating CRISPRi/CRISPRa functionality in a panel of multiple myeloma lines**

**A**. Multiple myeloma cell lines expressing CRISPRi machinery were transduced with control sgRNA or sgRNA targeting CD38. sgRNA-expressing cells were analyzed by flow cytometry for expression of CD38. Histograms indicate distribution of CD38 levels.
**B**. AMO1 cells expressing CRISPRa machinery were transduced with control sgRNA or sgRNA targeting CXCR4. sgRNA expressing cells were analyzed by flow cytometry for expression of CXCR4. Histograms indicate distribution of CXCR4 levels.

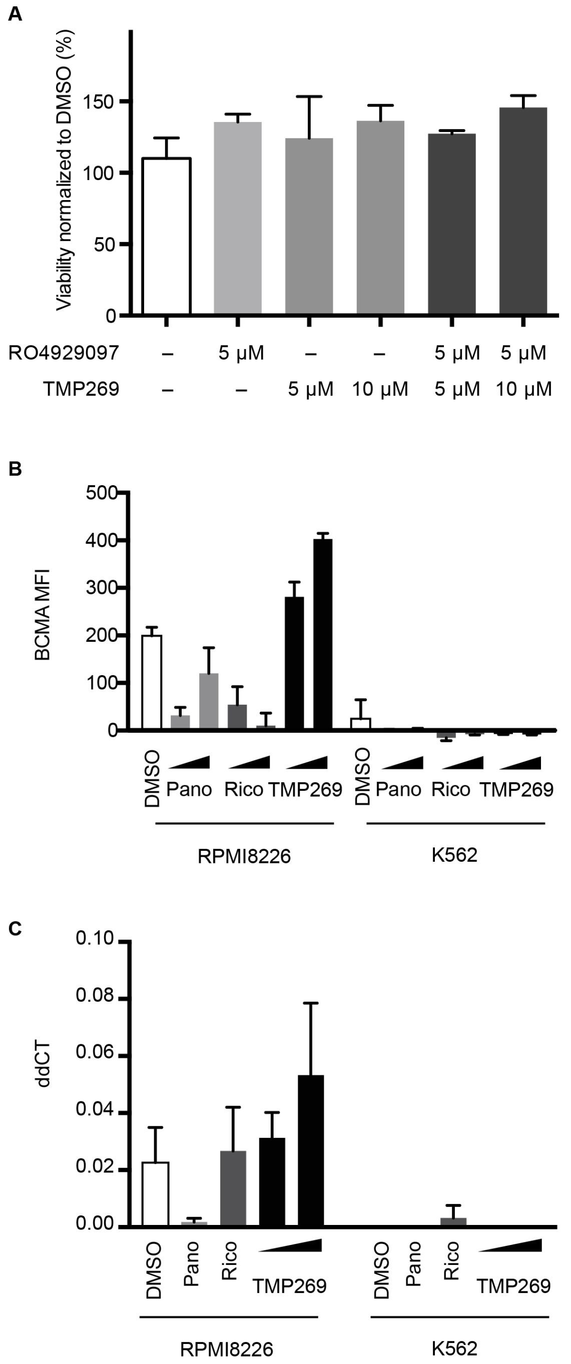


**Supplemental Fig 2:**
**Class IIa-HDAC inhibition is not toxic to MM cells and does not upregulate BCMA in K562 cells**

**A**. Viability of RPMI8226 cells post treatment with different drugs was determined by flow cytometry. Data indicates viability of cells normalized to DMSO control. Data points are means of three biological replicates; error bars denote SD.

**B**. RPMI8226 and K562 cell lines were treated with increasing concentrations of panobinostat (10 nM, 25 nM), ricolinostat (0.5 μM, 1 μM), TMP269 (5 μM and 10 μM) or DMSO for 48 hrs and analyzed by flow cytometry for cell-surface levels of BCMA. Bar graphs indicate the median fluorescence intensity (MFI) of BCMA after correcting for background fluorescence from isotype IgG control. Data points are means of three biological replicates; error bars denote SD.

**C**. RPMI8226 and K562 cell lines were treated with 10 nM Panobinostat, 0.5 μM Ricolinostat and 5 μM and 10 μM of TMP269 for 48 hrs were processed for quantitative PCR (qPCR) to determine BCMA transcript levels. Bar graphs represent delta delta CT (ddCT) values after normalizing to beta-actin gene. Data points are means of two biological replicates; error bars denote SD.

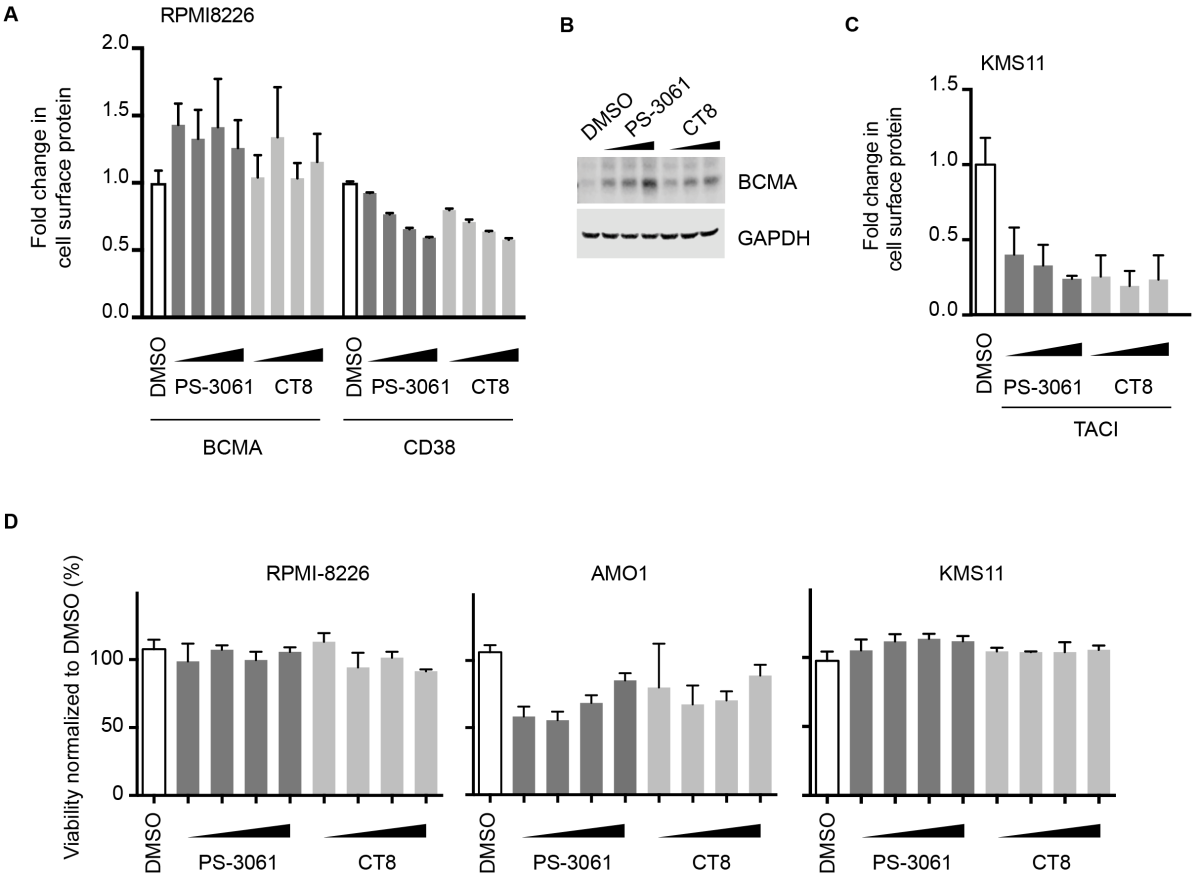


**Supplemental Fig 3:**
**Sec61 inhibition selectively increases levels of BCMA**

**A**. RPMI8226 cells were treated with increasing concentration of SEC61 inhibitors, PS3061 and CT8 (100, 200, 400, 800nM) or DMSO as a control for 24hrs. Cells were stained for cell surface expression of BCMA and CD38 and analyzed by flow cytometry. Data are means of three biological replicates; error bars denote SD.

**B**. Total protein lysates from RPMI8226 cells treated with increasing concentration of CT8 and PS3061 (200, 400, 800 nM) for 24 hrs were processed for Western blotting and immunoblotted for BCMA and GAPDH as the loading control.

**C**. KMS11 cells treated with increasing concentrations (200, 400, 800 nM) of SEC61 inhibitors, PS3061 and CT8, were stained for cell surface expression of TACI and analyzed by flow cytometry. Data are means of three biological replicates; error bars denote SD.

**D**. Viability of MM cells post treatment with different drugs was determined by flow cytometry. Data indicates viability of cells normalized to DMSO control. Data points are means of three biological replicates; error bars denote SD.

**A**

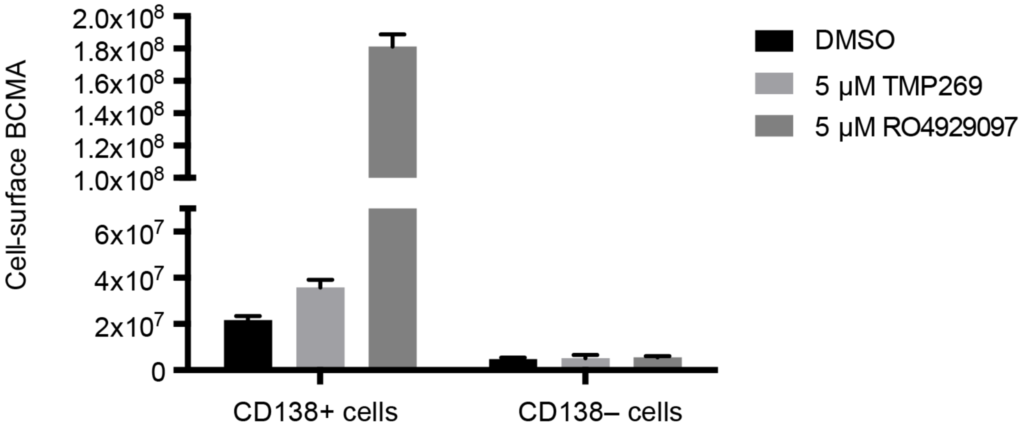


**B**

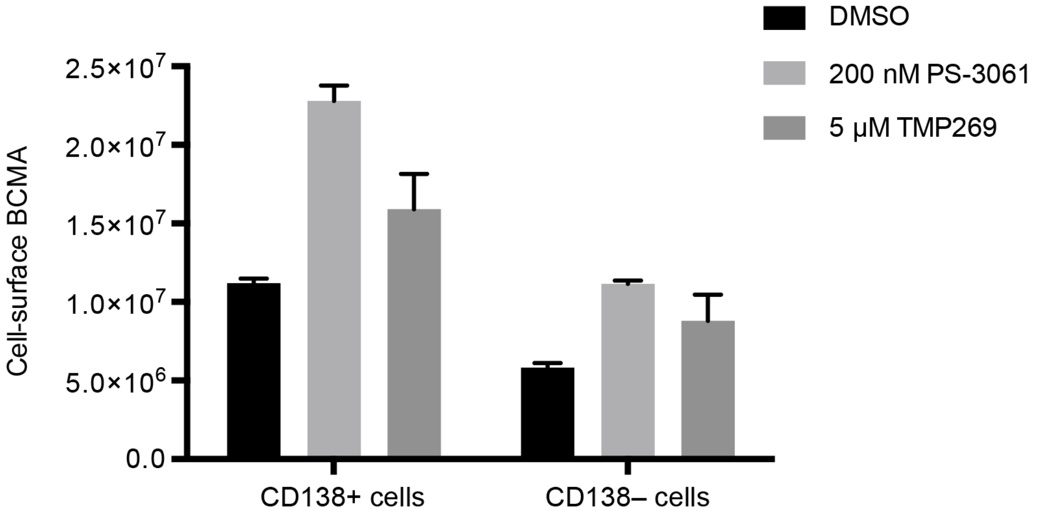


**Supplemental Fig. 4:**

**Pharmacological inhibition of the validated hits in myeloma patient samples show marginal effect on BCMA levels in non-plasma cells**. Bone marrow-mononuclear cells (BM-MNCs) isolated from bone marrow aspirates from different MM patients (A,B) were treated with indicated concentration of Class II-HDAC inhibitor, TMP269; gamma-secretase inhibitor, RO4929097; and Sec61 inhibitor, PS3061 for 24 hrs. Cells were stained for cell-surface CD138 and BCMA and analyzed by flow cytometry. BCMA Median Fluorescence Intensity was determined in both CD138-positive plasma cells and CD138-negative live cell populations. Data are means of three technical replicates, error bars denote SD.

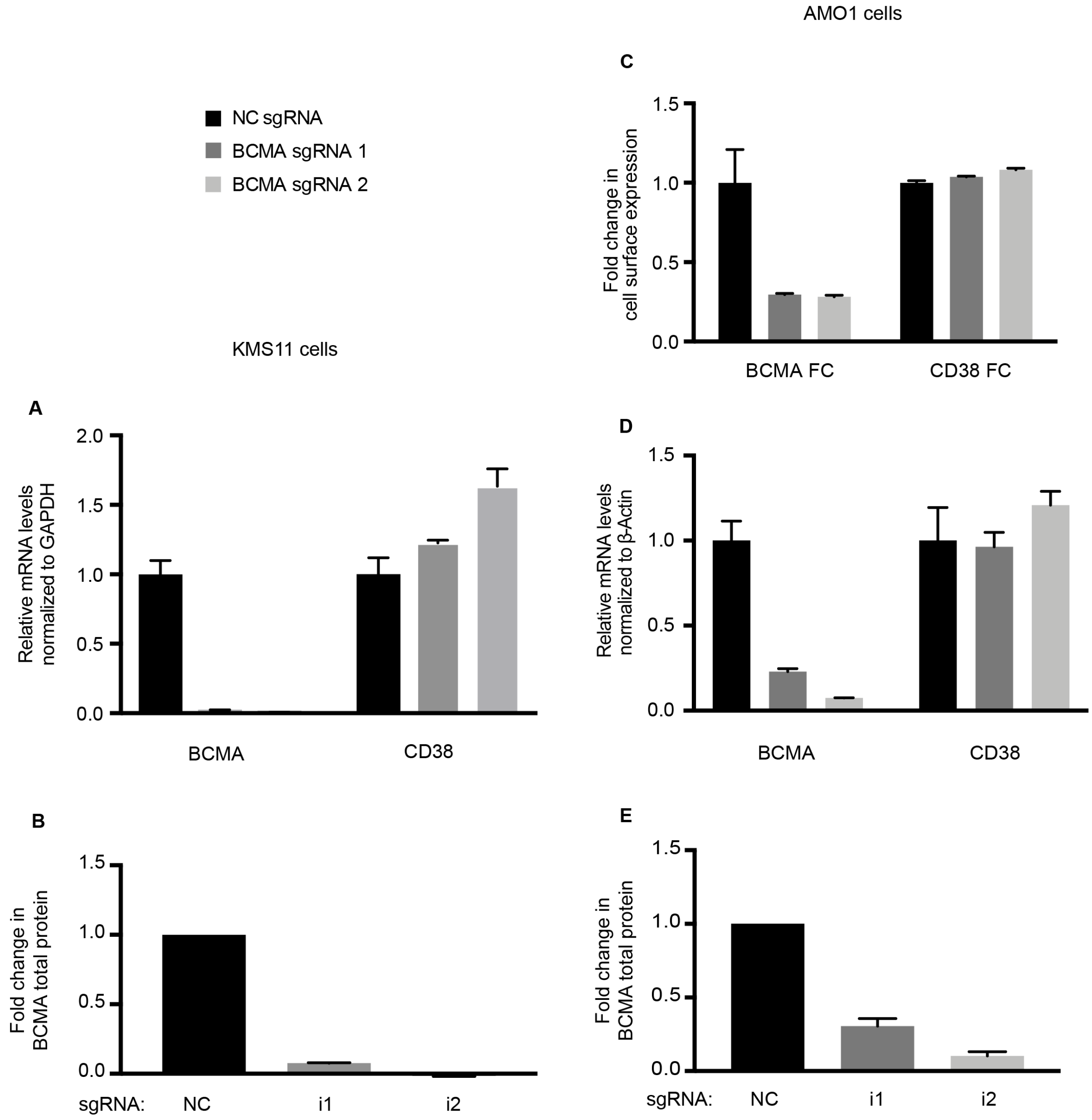


**Supplemental Fig. 5**

**Validation of BCMA knockdown in Myeloma cell lines**

**A,B,** KMS11 cells expressing control sgRNA and two independent sgRNAs targeting BCMA were: A. processed for qPCR to determine transcript levels of BCMA and CD38. Fold change in transcript levels were determined after normalizing to GAPDH. Data are means of two biological replicates; error bars denote SD. B. Total protein lysates from cells expressing the sgRNAs were processed for western blotting. GAPDH was used to normalize for differences in loading amounts. Data is represented as fold change relative to the normal protein expression level after normalization with GAPDH. Data points are means of two technical replicates, error bars denote SD.

**C-E**, AMO1 cells expressing control sgRNA and two independent sgRNAs targeting BCMA were: A. stained for cell surface BCMA and analyzed by flow cytometry. Data are means of three biological replicates, error bars denote SD., B. processed for qPCR to determine transcript levels of BCMA and CD38. Fold change in transcript levels were determined after normalizing to beta-actin. Data are means of two biological replicates; error bars denote SD. C. Total protein lysates from cells expressing the sgRNAs were processed for western blotting. GAPDH was used to normalize for differences in loading amounts. Data is represented as fold change relative to the normal protein expression level after normalization with GAPDH. Data points are means of two technical replicates, error bars denote SD.

Supplemental References


1. Lam C, Ferguson ID, Mariano MC, et al. Repurposing tofacitinib as an anti-myeloma therapeutic to reverse growth-promoting effects of the bone marrow microenvironment. *Haematologica*. 2018;103(7):1218-1228.

2. Kobro-Flatmoen A, Witter MP. Neuronal chemo-architecture of the entorhinal cortex: A comparative review. *Eur J Neurosci*. 2019;50(10):3627-3662.

3. Naumann RK, Ray S, Prokop S, Las L, Heppner FL, Brecht M. Conserved size and periodicity of pyramidal patches in layer 2 of medial/caudal entorhinal cortex. *J Comp Neurol*. 2016;524(4):783-806.

4. Mackinnon AL, Paavilainen VO, Sharma A, Hegde RS, Taunton J. An allosteric Sec61 inhibitor traps nascent transmembrane helices at the lateral gate. *Elife*. 2014;3:e01483.

5. Shah PS, Link N, Jang GM, et al. Comparative Flavivirus-Host Protein Interaction Mapping Reveals Mechanisms of Dengue and Zika Virus Pathogenesis. *Cell*. 2018;175(7):1931-1945 e1918.

6. Horlbeck MA, Gilbert LA, Villalta JE, et al. Compact and highly active next-generation libraries for CRISPR-mediated gene repression and activation. *Elife*. 2016;5.

7. Kampmann M, Bassik MC, Weissman JS. Functional genomics platform for pooled screening and generation of mammalian genetic interaction maps. *Nat Protoc*. 2014;9(8):1825-1847.

8. Chen JJ, Nathaniel DL, Raghavan P, et al. Compromised function of the ESCRT pathway promotes endolysosomal escape of tau seeds and propagation of tau aggregation. *J Biol Chem*. 2019.

9. Tak YE, Kleinstiver BP, Nunez JK, et al. Inducible and multiplex gene regulation using CRISPR-Cpf1-based transcription factors. *Nat Methods*. 2017;14(12):1163-1166.
